# DNA Replication Modulates R-loop Levels to Maintain Genome Stability

**DOI:** 10.1101/155978

**Authors:** Stephan Hamperl, Joshua Saldivar, Michael Bocek, Karlene A. Cimprich

## Abstract

Conflicts between transcription and replication are a potent source of DNA damage. The transcription machinery has the potential to aggravate such conflicts by generating co-transcriptional R-loops as an additional barrier to replication fork progression. Here, we investigate the influence of conflict orientation and R-loop formation on genome stability in human cells using a defined episomal system. This approach reveals that head-on (HO) and co-directional (CD) conflicts induce distinct DNA damage responses. Unexpectedly, the replisome acts as an orientation-dependent regulator of R-loop levels, reducing R-loops in the CD orientation but promoting their formation in the HO orientation. Replication stress and deregulated origin firing increase the number of HO collisions leading to genome-destabilizing R-loops. Our findings not only uncover an intrinsic function of the replisome in R-loop homeostasis, but also suggest a mechanistic basis for genome instability associated with deregulated DNA replication, which is observed in many disease states, including cancer.

## Introduction

During DNA synthesis, the replication machinery has to overcome numerous obstacles, including tightly bound DNA-protein complexes, non-B form DNA structures and DNA lesions that interfere with replication-fork progression (Mirkin and Mirkin, 2007). Failures to overcome these barriers lead to genome instability, a well-known hallmark of cancer and aging and a contributing factor to many other human disorders (Gaillard et al., 2015; Vijg and Suh, 2013; Zeman and Cimprich, 2014). Transcription complexes are one endogenous impediment frequently encountered by replication forks, and the resulting transcription-replication conflicts (TRCs) can induce DNA replication-fork stalling, DNA recombination, DNA breaks and mutations (Dutta et al., 2011; French, 1992; Liu and Alberts, 1995; Merrikh et al., 2011; Paul et al., 2013; Sankar et al., 2016; Srivatsan et al., 2010). Therefore, TRCs pose a potent threat to genome stability (García-Muse and Aguilera, 2016; Hamperl and Cimprich, 2016; Merrikh et al., 2012).

TRCs occur in two orientations: co-directional (CD), where the replication fork moves in the same direction as the transcription machinery, and head-on (HO), where the two converge on the DNA template. Bacterial studies suggest that HO collisions are the main cause of genomic alterations, leading to deletions, recombination and cell death (Srivatsan et al., 2010; Vilette et al., 1996). Thus, in bacterial genomes highly transcribed and essential genes are preferentially co-oriented with replication to minimize the number and consequences of HO-TRCs (Rocha, 2008). However, this orientation bias increases the frequency of CD collisions, and recent work indicates that they can disrupt replication at the highly transcribed ribosomal DNA clusters (Merrikh et al., 2011). These CD encounters can also cause double-strand break (DSB) formation on a plasmid when a stable backtracked RNA polymerase (RNAP) is encountered by the replication fork (Dutta et al., 2011).

In eukaryotic cells, TRCs have also been proposed to be a potent threat to genome stability. RNAP II transcribed genes can induce recombination when transcription and replication are oriented in opposite directions on yeast plasmid constructs (Prado and Aguilera, 2005). Furthermore, gene expression in mammalian cells can provoke S-phase-dependent recombination (Gottipati et al., 2008), and torsional stress created by depletion of Topoisomerase I leads to replication fork stalling and DNA breaks at certain S-phase transcribed genes (Tuduri et al., 2009). Collisions in long genes may be particularly difficult to avoid, because they undergo transcription for longer than one cell cycle. Moreover, some long genes overlap with common fragile sites, loci that replicate late in S-phase and represent hotspots for chromosomal instability (Helmrich et al., 2011; Le Tallec et al., 2014). Another class of fragile sites named early-replicating fragile sites are found at clusters of highly transcribed genes replicated in early S-phase (Barlow et al., 2013). Thus, it is tempting to speculate that their fragility is also driven by transcription-replication encounters. Together, these studies support the notion that conflicts between transcription and replication are frequent events in mammalian cells and can have detrimental effects on genome integrity.

One potent co-transcriptional block to the replication fork is the R-loop, an RNA-DNA hybrid which forms upon reannealing of a nascent transcript with the template DNA strand (Aguilera and García-Muse, 2012). Elevated levels of R-loops cause DNA damage and genome instability. The loss of RNA processing and R-loop regulatory factors increases R-loop levels and R-loop-dependent DNA damage in both yeast and human cells (Huertas and Aguilera, 2003; Li and Manley, 2005; Paulsen et al., 2009; Santos-Pereira and Aguilera, 2015; Sollier et al., 2014). This R-loop-induced DNA damage is dependent upon DNA replication, consistent with the finding that some of these R-loops act as a barrier for the replication fork (Castellano-Pozo et al., 2012; Gan et al., 2011; Tuduri et al., 2009; Wellinger et al., 2006). However, the impact of collision orientation on this process and the molecular steps leading to the activation of DNA damage signaling pathways and genome instability are unclear. Intriguingly, R-loops are also prevalent in eukaryotic genomes (Sanz et al., 2016) and are involved in diverse cellular processes, including immunoglobulin class switch recombination (Yu et al., 2003), transcription termination (Skourti-Stathaki et al., 2011), and the regulation of gene expression (Sun et al., 2013). If and how these regulatory R-loops are tolerated or removed during DNA replication is unknown.

The plasticity of origin firing during eukaryotic DNA replication poses several challenges to studying TRCs, the impact of R-loops on TRCs, and the mechanisms by which they cause DNA damage in mammalian cells. Unlike in bacteria, where a single origin allows for replication of each gene in a predictable fashion, mammalian chromosomes contain a multitude of replication origins (Hills and Diffley, 2014). These fire with variable efficiencies and timing (McGuffee et al., 2013; Rhind and Gilbert, 2013). Moreover, there is an excess of origins that can be used to complete DNA synthesis from the opposing direction when a replication fork stalls (Alver et al., 2014). Thus, it is difficult to predict the location and orientation of a TRC in a population of cells.

Here, we report a novel system that allows for exquisite control over the direction and timing of replication, transcription and R-loop formation on plasmids stably maintained in human cells. We observe striking differences between HO and CD collisions in the context of R-loop formation. These different collisions induce distinct DNA damage responses and differentially affect the stability of R-loops, reducing R-loops in the co-directional orientation but promoting their formation in the head-on orientation. Importantly, we demonstrate the same effects in the native genomic context. These observations suggest that the replisome is an orientation-dependent regulator of R-loop levels and provide mechanistic insight into how regulatory R-loops are tolerated in S phase. They also indicate that R-loop levels may be perturbed under replication stress and may represent the underlying source of genomic instability in cells that experience replication stress, a hallmark of cancer cells and many other disease states (Gaillard et al., 2015; Vijg and Suh, 2013; Zeman and Cimprich, 2014).

## Results

### An episomal system to study transcription-replication conflicts

To systematically probe the events that occur upon TRCs, we developed an episomal system to induce these conflicts in a localized and controlled fashion. We took advantage of the Epstein–Barr virus replication origin (oriP) which allows for unidirectional replication of chromatinized plasmids ((Kirchmaier and Sugden, 1995) and data not shown). The co-expressed Epstein–Barr virus nuclear antigen 1 (EBNA1) protein utilizes the endogenous replication machinery, including the MCM replicative helicase, but it also blocks progression of one arm of the bidirectional replication fork by creating a replication fork barrier at the direct repeats of oriP (Dhar et al., 2001; Gahn and Schildkraut, 1989). We cloned the R-loop forming portion of the mouse AIRN (mAIRN) gene (Ginno et al., 2012) or the non R-loop forming ECFP sequence in both directions relative to oriP to compare R-loop dependent (mAIRN) and independent (ECFP) HO and CD collisions (Figure 1A). The formation of R-loops at both sequences independent of replication was analyzed by *in vitro* transcription of bacterial plasmid templates. Stable RNA-DNA hybrids induce a topological change visualized by reduced mobility in native agarose gel electrophoresis (Ginno et al., 2012). As expected, transcription of the mAIRN sequence, but not of the ECFP control sequence, resulted in extensive, RNase H-sensitive formation of RNA-DNA hybrids *in vitro* (Figure 1B).

**Figure 1.**
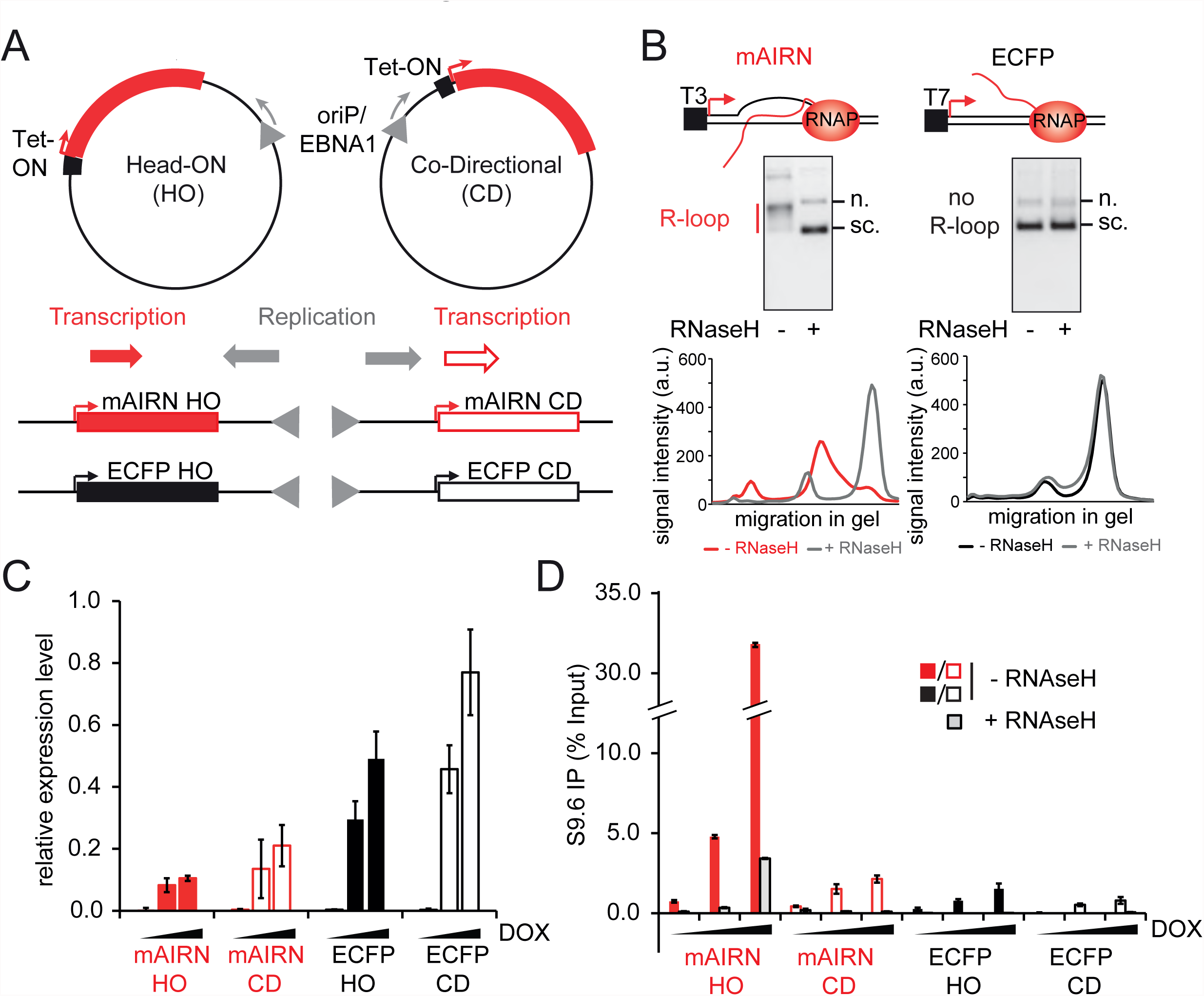
An episomal system to study transcription-replication conflicts. A) Schematic representation of the constructs. The mAIRN and ECFP transcription units were placed in a HO or CD orientation with the oriP origin of replication derived from the Epstein-Barr virus. B) R-loop formation at the mAIRN sequence *in vitro*. Plasmids containing the mAIRN or ECFP sequence were transcribed *in vitro*. Samples were split equally and treated, or not, with RNase H (indicated by + or -). The upper panel is the gel after ethidium bromide staining, while the bottom panel corresponds to a profile analysis of individual gel lanes. Signal intensities were plotted against the migration in the gel (a.u. arbitrary units). C) RT-qPCR analysis of mAIRN and ECFP HO/CD-induced transcription. RNA samples were extracted from cells 72h after treatment with 0, 100 or 1000 ng/mL DOX. Gene expression was normalized relative to the expression of the β-actin genomic locus. The bars indicate mean and standard deviations between biological replicates (n=3). D) DRIP-qPCR analysis of mAIRN and ECFP HO and CD constructs. Cells were treated with 0, 100 or 1000 ng/mL doxycycline in the culture medium for 72h and harvested for DRIP. The bars indicate mean and standard deviations between biological replicates (n=2).

Next, we introduced the oriP-containing constructs into human embryonic kidney (HEK293) cells that express the tetracycline (Tet)-regulated transactivator, allowing for doxycycline (DOX)-induced transcription of the Tet-ON promoter-controlled mAIRN and ECFP sequences. After transfection, we generated stable cell lines that maintain 10 to 150 plasmid copies per cell with >90% efficiency per generation (Figure S1A-C). This observed rate of plasmid loss is in good agreement with previous studies that used similar oriP/EBNA1-based vectors (Leight and Sugden, 2001). Reverse-transcription quantitative PCR (RT-qPCR) revealed dose-dependent DOX-induced transcriptional activation of all Tet-ON controlled sequences in all cell lines (Figure S2A-D). Interestingly, transcription of the mAIRN sequence was impaired when placed in the HO versus CD orientation, and a similar ~5-fold orientation-dependent difference was observed with the ECFP constructs. These findings suggest that head-on gene expression has an inhibitory effect on RNAP II transcription (Figure 1C and Figure S2). Independent of gene orientation, we also noticed that overall DOX-induced transcription was significantly higher with the ECFP versus R-loop forming mAIRN sequences (Figure 1C and Figure S2). Although we cannot exclude that this difference in the steady-state transcription levels is caused by differences in mRNA stability, these data imply that gene expression is also impaired by the presence of R-loops on the mAIRN transcription unit.

To determine whether transcription induces R-loop formation on the mAIRN sequence in cells, we performed DNA-RNA immunoprecipitation (DRIP) and qPCR on the plasmid using the RNA-DNA hybrid-specific S9.6 antibody (Boguslawski et al., 1986). We found that the mAIRN HO cells exhibited robust RNase H-sensitive RNA-DNA hybrid formation after DOX addition (Figure 1D, mAIRN HO). Surprisingly, however, RNA-DNA hybrids were poorly induced in the mAIRN CD cells and found at levels only slightly above those observed in the ECFP HO and CD control cells (Figure 1D). Thus, RNA-DNA hybrids were specifically enriched in the mAIRN HO cells, despite the fact that the same sequence under control of the same promoter was transcribed in mAIRN CD cells. This finding suggests that the orientation with which a replication fork approaches the mAIRN transcription unit critically affects hybrid formation or hybrid stability on the plasmid.

### The orientation of transcription and replication affects RNA-DNA hybrid levels

To further investigate the distinct effects of HO and CD TRCs on hybrid levels, we took advantage of the ability to synchronize plasmid replication and analyzed hybrid formation on the mAIRN HO and CD constructs before and after release from a single-thymidine block (Figure 2A and Figure S3A-B). We reasoned that if this difference in hybrid levels is dependent on DNA replication, arresting cells in G1/S should alleviate this difference by preventing DNA replication on the constructs. Successful arrest at the G1/S transition, release into S-phase and transcriptional induction were confirmed by flow cytometry and RT-qPCR analyses (Figure S3A-C). As we previously observed, hybrid formation was more robust on the mAIRN HO plasmid than on the mAIRN CD plasmid in asynchronous cells (Figure 2B, ASYN). However, arrest of cells at the G1/S transition reduced hybrids on the mAIRN HO plasmid and increased hybrids on the mAIRN CD plasmid. Strikingly, this block to replication neutralized the orientation-dependent difference in hybrid levels (Figure 2B, G1/S). Moreover, release of cells into S-phase restored the difference (Figure 2B, S). These observations provide strong evidence that replication fork progression increases co-transcriptional hybrid levels on the mAIRN HO construct, whereas it decreases hybrid levels on the CD construct. Thus, we conclude that the replisome is an orientation-dependent regulator of RNA-DNA hybrid levels in our episomal system.

**Figure 2.**
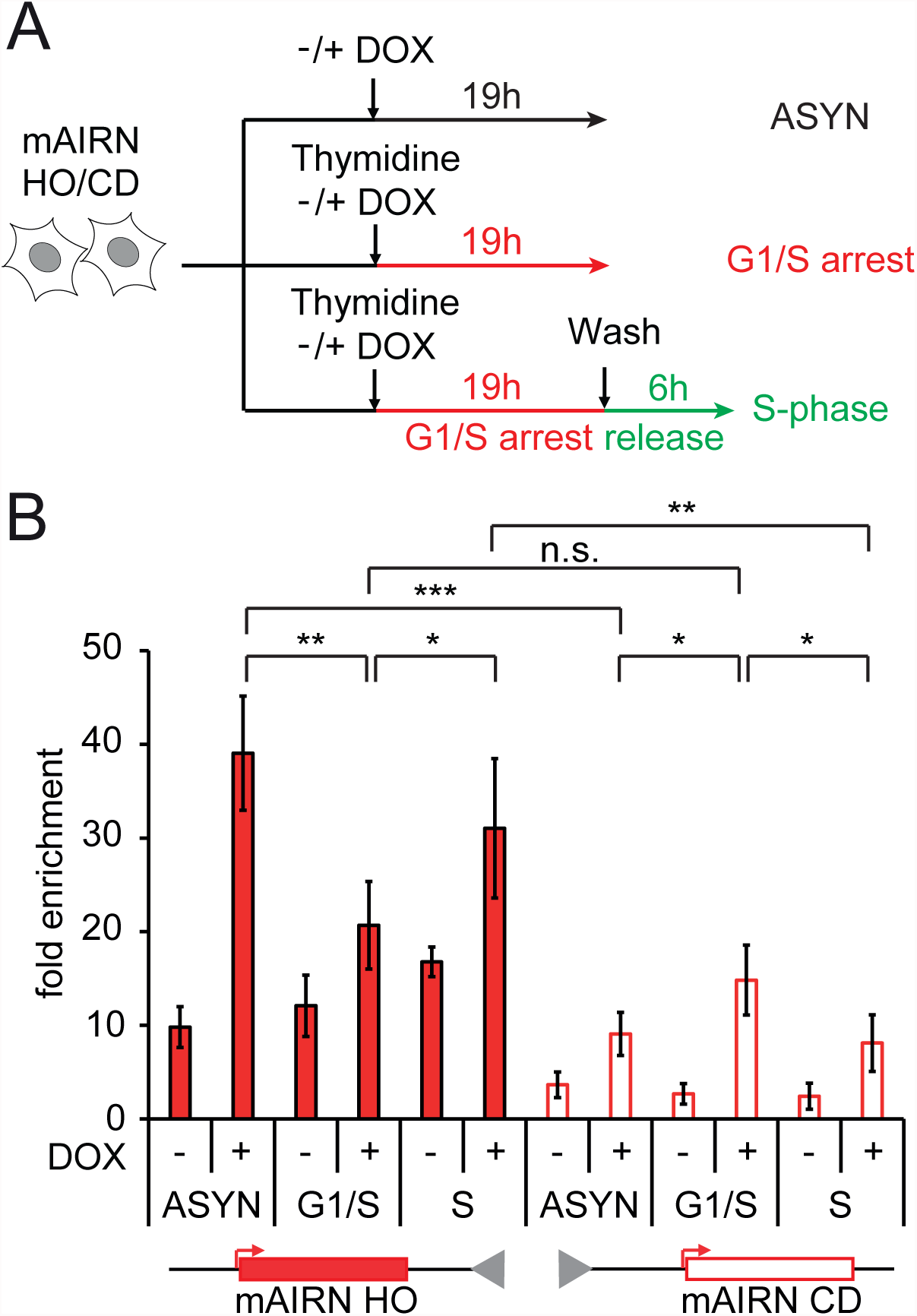
The orientation of transcription and replication affects RNA-DNA hybrid levels. A) Schematic for the single-thymidine block and release experiment. B) DRIP-qPCR analysis of mAIRN HO and CD cells after G1/S arrest or release into S-phase either without (-) or with (+) addition of 1000 ng/mL DOX. As a control, asynchronously growing cells without (-) or with (+) 1000 ng/mL DOX for 19h were simultaneously harvested for DRIP (ASYN). The DRIP signals were normalized and shown as fold enrichment to the non R-loop forming negative control ZNF544 locus on the genome. The bars indicate mean and standard deviations between biological replicates (n=4). \**p*<0.05. **p<0.01. ***p<0.001, unpaired Student’s t-test.

### R-loop formation exacerbates the effect of TRCs on plasmid instability

We next tested the effects of HO and CD collisions on plasmid stability. At the highest concentration of DOX, the mAIRN HO plasmid copy number was reduced to ~27% of that observed without DOX stimulation, while the copy number of the mAIRN CD plasmid was reduced to ~50% (Figure 3A). This effect was only observed after prolonged transcriptional induction (≥ 24h), suggesting that cell cycle progression is required to induce significant plasmid loss (Figure S4A-B). Increased transcription-induced plasmid loss in mAIRN HO versus mAIRN CD cells was also confirmed by Southern blot analysis of extracted genomic DNA probed against the plasmid or a region of the genomic β-actin gene (Figure S4C-D). The ECFP sequence showed a similar orientation-dependent difference in plasmid stability (Figure 3A, ~42% HO and ~83% CD). However, plasmid loss was more pronounced with the R-loop prone mAIRN sequence, even at lower transcription levels. In order to provide more direct evidence that plasmid instability is exacerbated by the presence of R-loops on the mAIRN sequence, we transfected the mAIRN and ECFP cell lines with a control vector or a vector overexpressing RNaseH1 and determined plasmid loss after activation of transcription (Figure 3B). Strikingly, plasmid instability was partially rescued in mAIRN HO cells, whereas no significant effect was observed in mAIRN CD cells or the ECFP control constructs. This is consistent with our previous finding that R-loop formation is most prominent in the mAIRN HO cells (Figure 1D) and suggests that a significant portion of the plasmid loss in this orientation is R-loop dependent. Together, these results indicate that HO collisions are more detrimental than CD collisions, regardless of R-loop formation. However, R-loop formation exacerbates the effect of TRCs on plasmid instability in both orientations.

**Figure 3.**
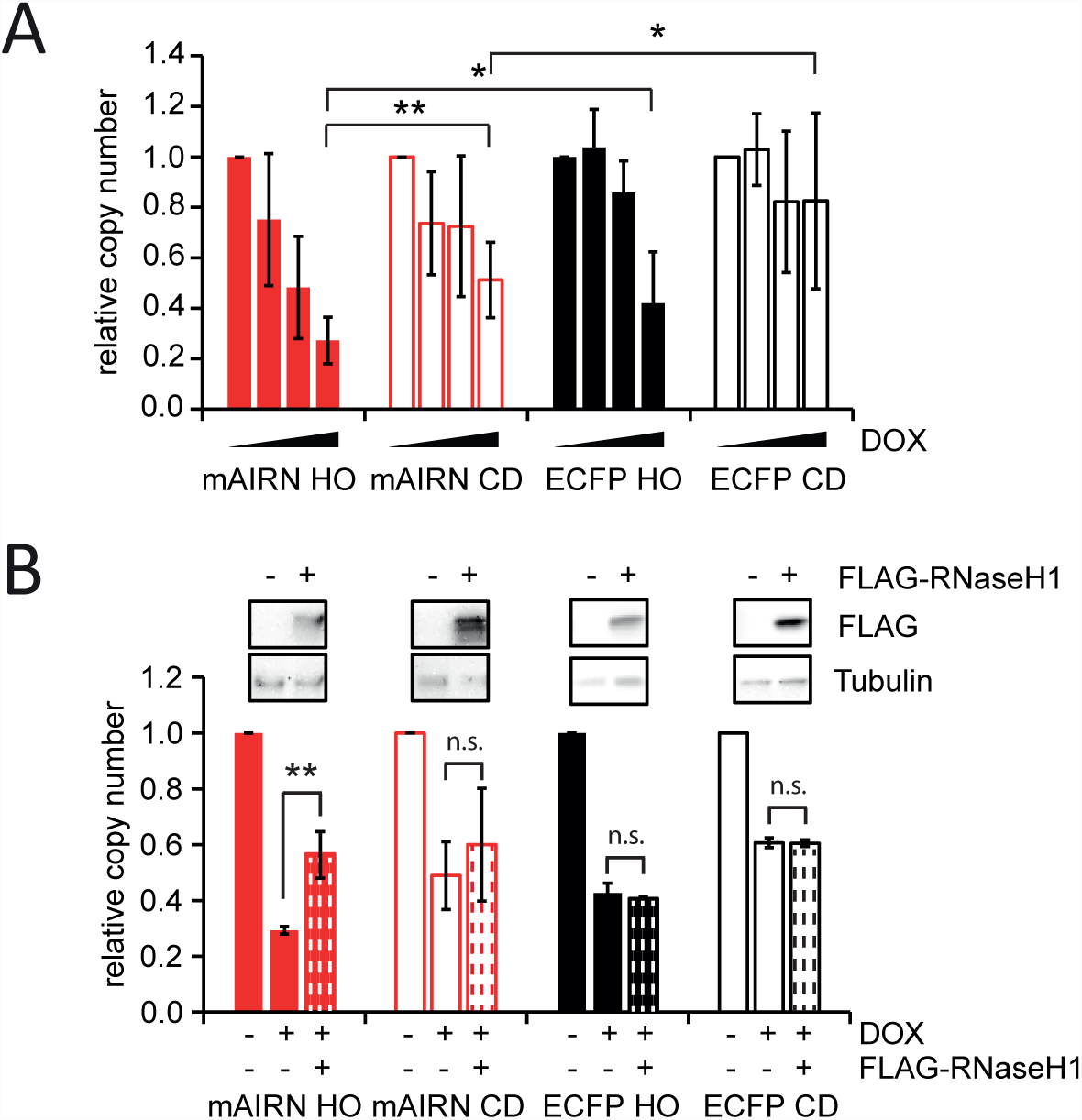
R-loop formation exacerbates the effect of TRCs on plasmid instability. A) mAIRN/ECFP HO and CD cells were treated with 0, 50, 100, or 1000 ng/mL DOX for 72h. After extraction of genomic DNA, the relative plasmid copy number (normalized to 0 ng/mL DOX) was determined by quantitative PCR. The bars indicate mean and standard deviations between biological replicates (n 3). *p<0.05. **p<0.01, unpaired Student’s t-test. B) mAIRN and ECFP HO and CD cells were treated with 0 (-) or 1000 ng/mL DOX (+). After 24h, cells were transfected with a vector expressing FLAG-tagged RNaseH1 (+) or an empty vector control (-) in the absence (-) or presence (+) of 1000 ng/mL DOX for further 48h. Plasmid copy number was determined as described above. The bars indicate mean and standard deviations between replicate experiments (n=2-4). ** p< 0.01, n.s. not significant, unpaired Student’s t-test.

### HO and CD conflicts on the mAIRN transcription unit induce distinct DNA damage responses

Next, we sought to determine whether transcription-induced plasmid instability is associated with DNA damage arising in response to TRCs on the R-loop forming constructs. To do so, we monitored activation of the cellular DNA damage response, first assessing the phosphorylation of H2AX (γ-H2AX) in these cells. Interestingly, γ-H2AX was induced in the mAIRN HO cells, but not in the mAIRN CD cells, despite the fact that the mAIRN CD cells carried ~5-fold more plasmids than the mAIRN HO cells (50-100 vs ~10-20 copies per cell) (Figure 4A). We did detect γ-H2AX induction in an mAIRN CD cell line with an even greater copy number (~140 copies) (Figure S5A), suggesting that a greater number of CD-TRCs is required to induce a detectable γ-H2AX response.

**Figure 4.**
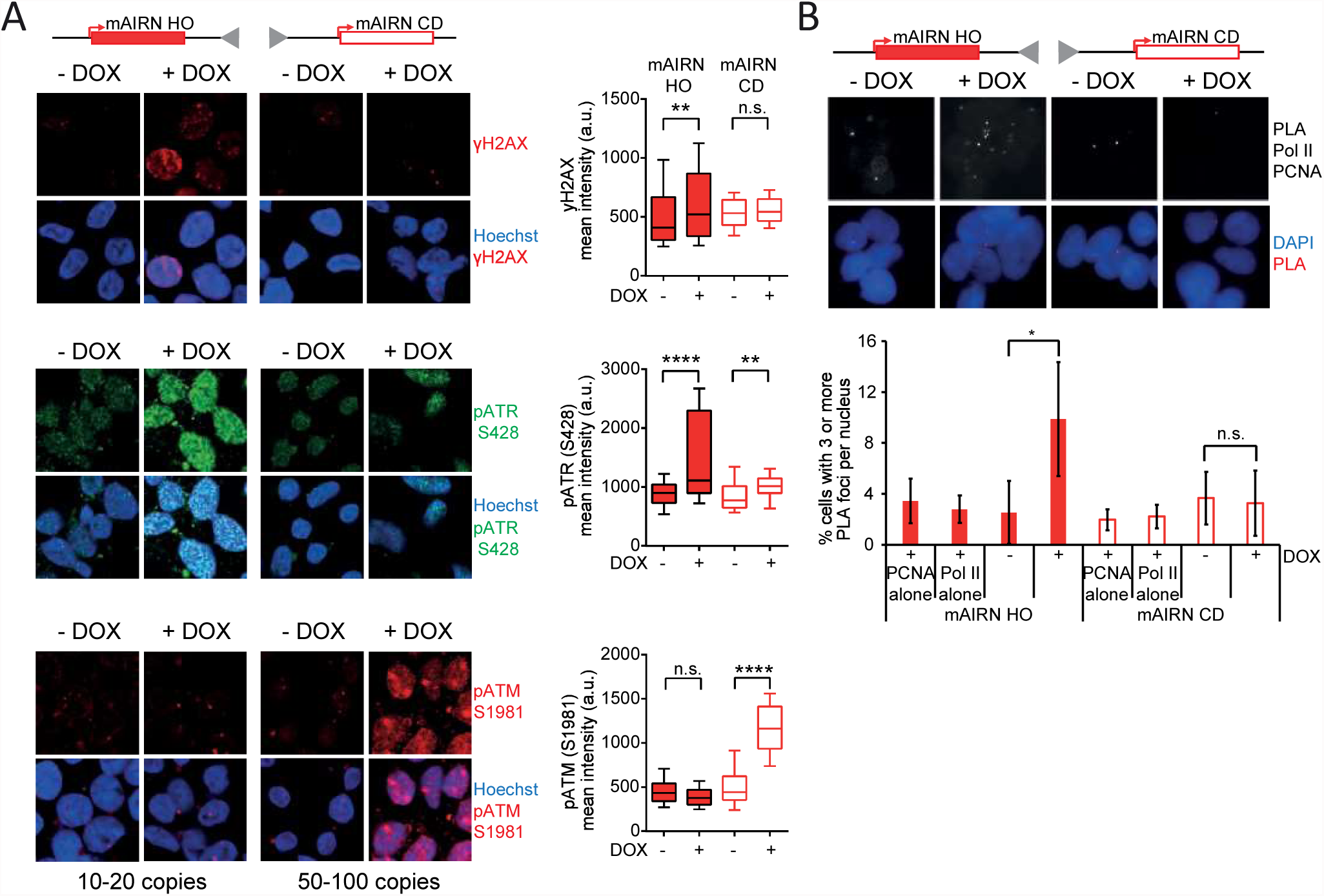
HO and CD conflicts induce distinct DNA damage responses. A) Representative images (left) and quantification (right) after immunostaining for γH2AX, ATR pS428 or ATM pS1981 in mAIRN HO and CD cells either treated with 0 or 1000 ng/mL DOX for 48h. Hoechst is used to stain the nucleus. Box and whisker plots show the 10-90 percentile. a.u. = arbitrary units. n.s. not significant. \*\**p*<0.01. \*\*\*\**p*<0.0001. One-way ANOVA test (n 100). B) HO conflicts result in persistent proximity between transcription and replication machineries. Representative images (top) and quantification (bottom) of the percentage of cells with 3 RNAP II-PCNA PLA foci per nucleus. DAPI is used to stain the nucleus. RNAP II alone and PCNA alone are single-antibody controls from mAIRN HO and CD cells treated with 1000 ng/mL DOX for 24 hours. The bars indicate mean and standard deviations between biological replicates (n ≥ 3). n.s. not significant. *p<0.05. Unpaired Student’s t-test.

As γ-H2AX was induced with distinct thresholds in mAIRN HO and CD cells, we considered the possibility that different DNA damage signaling pathways are activated. Thus, we monitored activation of the DNA damage response kinases ATR (ataxia telangiectasia-mutated [ATM] and rad3-related) and ATM. Whereas ATM is primarily activated by DSBs, ATR responds to stretches of RPA-coated single-stranded DNA (ssDNA) originating from events that cause replication fork stalling (Cimprich and Cortez, 2008; Maréchal and Zou, 2013). Transcriptional activation of the mAIRN HO construct induced ATR S428 phosphorylation, whereas ATM S1981 autophosphorylation was unaffected. In contrast, transcriptional activation in the mAIRN CD cells resulted in strong ATM autophosphorylation, but had only a minor effect on ATR phosphorylation (Figure 4A and Figure S5B-C). Importantly, we also observed preferential phosphorylation of other ATM targets including Chk2 and KAP1 S824 in mAIRN CD cells, whereas the ATR effector kinase Chk1 and the ATR target RPA2 S33 were preferentially phosphorylated in mAIRN HO cells (Figure S6A-B). Taken together, our findings demonstrate that HO and CD TRCs induce distinct DNA damage responses, likely through the formation of distinct structures, with the HO collision giving rise to ATR activation and the CD collision leading to ATM activation.

ATR activation in mAIRN HO cells suggests that replication forks stall at transcription complexes in this orientation. To more directly test this idea, we sought to detect interactions between the transcription and replication machineries using the proximity-ligation assay (PLA). We used antibodies against RNAP II as a marker for transcription complexes and PCNA as a marker of replication forks. In pre-extracted cells containing the mAIRN HO plasmid, we detected a ~4-fold increase in the percentage of cells that contain ≥3 PLA foci (RNAP II and PCNA interaction) upon DOX induction. Strikingly, no change was detected in the CD orientation with this assay (Figure 4B). This finding suggests that persistent collisions arise from HO but not CD TRCs, consistent with activation of ATR.

### R-loops are enriched at HO regions of the genome

In order to test the generality of our conclusions from the plasmid system, we asked if the orientation and frequency of TRCs modulates RNA-DNA hybrid levels in the native genomic context. For this approach, we examined the distribution of RNA-DNA hybrids around replication origins in the genome. To do so, we performed DRIP followed by next generation sequencing (DRIP-Seq) to determine the location and relative abundance of R-loops genome-wide in unperturbed HeLa cells. Next, we used a published Okazaki fragment sequencing (OK-Seq) dataset from the same cell line to identify the genomic locations of strong replication origins (Petryk et al., 2016). We then identified the subset of these origins that are proximal to transcribed units using publically available HeLa Global Run-On Sequencing (GRO-Seq) data (Andersson et al., 2014). Using this dataset to additionally define the direction of transcription around the origin, we identified specific subsets of origins that are in close proximity to transcription units on both sides. In this way, we called 1,084 origin-proximal regions with HO transcription on either side (HO-HO), 1,025 regions with CD transcription (CD-CD), and 6,703 regions with HO transcription on one side and CD transcription on the other (HO-CD) (Figure 5A-B). We then examined DRIP-Seq read density in a 30kb window around each class of origin to determine the relative levels of hybrids in each situation (Figure 5C). Finally, we compared the levels of DRIP signal between these classes, accounting for systematic differences in factors known to correlate with hybrid formation, including total transcription levels, replication timing or GC-content (see bioinformatic and statistical analyses for details). Strikingly, origin-proximal regions that were classified as HO-HO showed a significant increase in DRIP signal relative to CD-CD regions on either side of the origin (Figure 5D). Moreover, HO-CD regions showed a similar DRIP enrichment at the HO side, but not at the side biased towards CD collisions. Thus, regions predicted to undergo HO collisions in the genome have a higher frequency of RNA-DNA hybrids than regions predicted to undergo CD collisions. These findings are consistent with data from our episomal system and our model that co-orientation of transcription and replication reduces origin-proximal R-loop levels.

**Figure 5.**
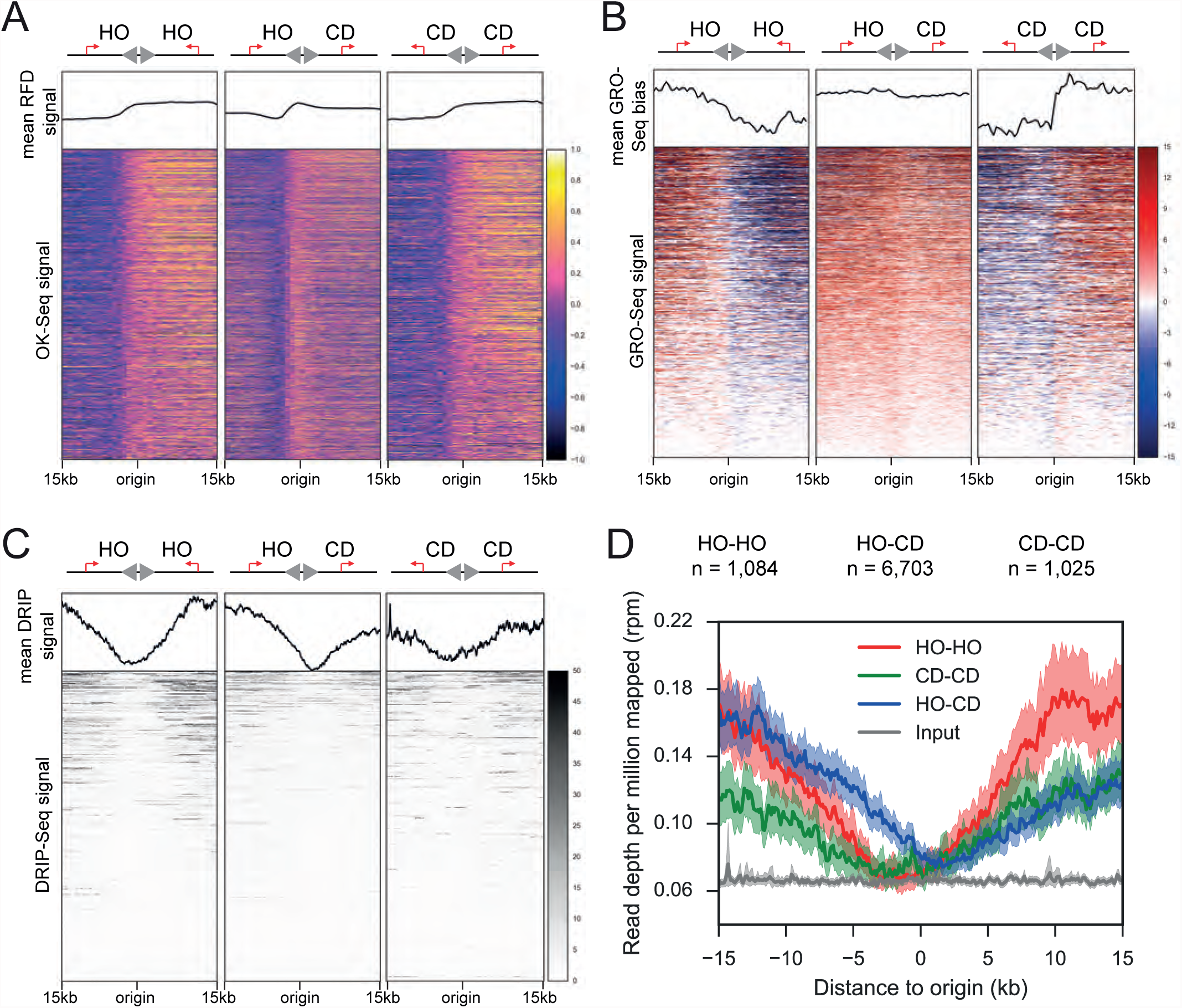
R-loops are enriched at HO regions of the genome. A-C) Heat maps of A) replication fork directionality (RFD) from OK-Seq data, B) GROSeq strand bias and C) DRIP-Seq in HeLa cells along the identified 1,084 HO-HO, 6,703 HO-CD and 1,025 CD-CD origin-proximal regions, centered where RFD crosses zero and sorted from highest to lowest total GRO-Seq reads from the positive and negative strands. The top panels of each heat map represent A) the aggregate mean RFD, B) GRO-seq strand bias and C) DRIP-Seq signals centered around the origins of each category. Origins were defined as locations where the RFD signal (A) changes sign from negative to positive, then categorized as HO-HO, CD-CD or HO-CD by the relative direction of transcription as found from the strandedness bias of GRO-seq reads (B.) D) Aggregate plot of DRIP signal (reads per million mapped) in HeLa cells centered around origins with a transcription bias towards HO-HO (red), HO-CD (blue) or CD-CD (green) orientation. Shaded areas represent 95% confidence intervals by a bootstrap of the mean.

### Perturbation of the replication program increases genomic levels of R-loops

Recent analysis of high resolution OK-Seq data indicates that the movement of replication forks and transcription complexes is significantly co-oriented in the human genome (Petryk et al., 2016), similar to what has been reported in bacterial genomes (Rocha, 2008). One prediction of our results is that this type of co-orientation bias would help to minimize genomic R-loops in S phase, whereas disruption of this bias would augment the genomic level of these structures. To test this hypothesis, we treated cells with high doses of hydroxyurea (HU) or the DNA polymerase inhibitor aphidicolin (APH) in order to abolish DNA synthesis. Blocked replication forks should be unable to traverse transcribed regions under these conditions, preventing hybrid clearance in the more common CD orientation and reducing the number of TRCs in the genome (Figure S7A-B). Indeed, we observed an increase in nuclear RNA-DNA hybrids after HU or APH treatment by S9.6 immunofluorescence (Figure 6A) and only background levels of TRCs as measured by RNAP II-PCNA PLA foci (Figure 6B). This result suggests that R-loop levels are reduced by active replication forks under normal conditions, and is consistent with a bias for co-directional organization of the human genome.

**Figure 6.**
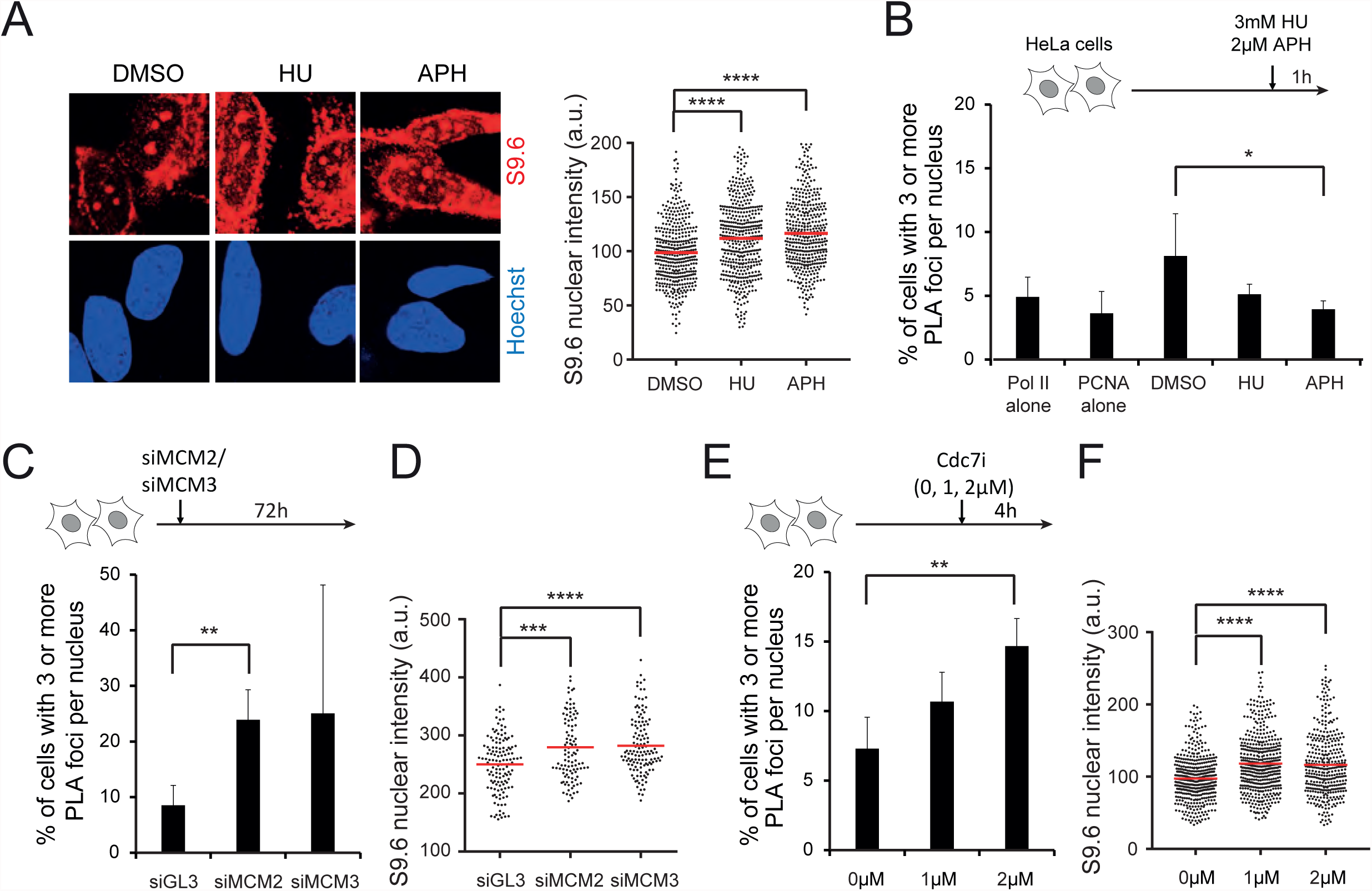
Perturbation of the replication program increases genomic levels of R-loops. A) Immunostaining (left) and quantification (right) of S9.6 nuclear signal in HeLa cells treated with 3mM HU or 2μM aphidicolin for 1h. The nucleus was co-stained with Hoechst. The mean value is shown as a red line. a.u. = arbitrary units. ****p<0.0001. One-way ANOVA test (n≥400). B) Percentage of cells with ≥ 3 RNA RNAP II and PCNA PLA foci under the same conditions as in A). RNAP II alone and PCNA alone are single-antibody controls from HeLa cells treated with DMSO for 1h. The bars indicate mean and standard deviations between biological replicates (n ≥ 3). *p<0.05. Unpaired Student’s t-test. C) Percentage of cells with ≥ 3 PLA foci between RNAP II antibody and PCNA antibody in HeLa cells transfected with indicated siRNAs and fixed after 72h. The bars indicate mean and standard deviations between biological replicates (n ≥ 3). **p<0.01. Unpaired Student’s t-test. D) Immunostaining and quantification of S9.6 nuclear signal in HeLa cells under the same conditions as in C). The nucleus was co-stained with Hoechst. The mean value is shown as a red line. a.u. = arbitrary units. ***p<0.001. ****p<0.0001. One-way ANOVA test (n≥100). E) Percentage of cells with ≥ 3 PLA foci between RNAP II antibody and PCNA antibody in HeLa cells treated with Cdc7 inhibitor (PHA-767491) at the indicated concentrations and fixed 4 hr later. The bars indicate mean and standard deviations between biological replicates (n ≥ 3). **p<0.01. Unpaired Student’s t-test. F) Immunostaining and quantification of S9.6 nuclear signal in HeLa cells under the same conditions as in E). The nucleus was co-stained with Hoechst. The mean value is shown as a red line. a.u. = arbitrary units. ****p<0.0001. One-way ANOVA test (n≥370).

Next, we sought to reverse this co-orientation bias and increase the number of HO collisions in the genome, a perturbation our model predicted to result in higher genomic R-loop levels. To do so, we monitored the effect of partially depleting Mcm2-7 complexes or inhibiting the Cdc7 kinase on RNA-DNA hybrid levels and RNAP II-PCNA PLA interactions. Mcm2-7 complexes are loaded in excess over the number of origins normally used, licensing dormant or backup origins that can rescue forks stalled at lesions and other barriers and allowing completion of DNA synthesis under conditions of replication stress (Ge et al., 2007; Ibarra et al., 2008). Cdc7 is an essential kinase that activates origins by phosphorylating the Mcm2-7 complex at the time of replication initiation (Jiang et al., 1999; Montagnoli et al., 2008). Although both perturbations target different steps of the replication cycle, both change the pattern of origin usage during DNA synthesis (Kunnev et al., 2015). This should lead some forks to travel for a longer distance and/or to approach transcription complexes with a different orientation, increasing the frequency of collisions in the undesirable HO orientation (Figure S7C) (Hills and Diffley, 2014). Consistent with this hypothesis, small interfering (si)RNA-mediated knockdown of Mcm2 or Mcm3, as well as Cdc7 kinase inhibition, resulted in a significant increase in the percentage of cells that contain ≥3 RNAP II-PCNA PLA foci (Figure 6C, E). As predicted, these perturbations also increased nuclear RNA-DNA hybrids (Figure 6D, F). Similar results were observed in a different cell line and with independent siRNAs (Figure S8A-C), and there was no effect of partial Mcm2 or Mcm3 depletion on cell cycle progression (Figure S8D). In addition, the doses used for the Cdc7 inhibitor did not significantly inhibit the Cdk9 kinase, which is involved in transcriptional regulation (Shim et al., 2002) (Figure S8E). These findings suggest that HO collisions and R-loops are induced by changes in origin firing preferences, and taken together with previous results provide strong evidence that DNA replication fork progression modulates R-loop formation in an orientation-dependent manner.

## Discussion

Using the unidirectional oriP/EBNA1 replicon in combination with different Tet-ON promoter controlled transcription units, we established an *in vivo* system to analyze encounters between the replication fork and different types of transcriptional barriers in an inducible and localized fashion. This system allows us to reliably define collision orientation, which is difficult on mammalian chromosomes due to their multiple and variable origins of replication. Thus, we could compare the destabilizing effects of HO versus CD-oriented collisions between a replication fork and co-transcriptional R-loops in human cells. Conflict orientation affected both the stability of the plasmid and the specific damage response pathway that was activated by the TRC (Figure 7), indicating that different ATR- or ATM-activating structures form depending on how the fork encounters these barriers. Strikingly, R-loop formation is also highly dependent on the orientation of the TRC. HO collisions increase R-loop formation, whereas CD collisions decrease R-loop formation. These findings suggest that the replisome can resolve R-loops in the CD orientation, demonstrating a new intrinsic function for this machinery. Overall, our studies provide mechanistic insights into how the threat of R-loops to genome stability during replication is balanced with their physiological roles through this function of the replisome, the organization of the genome and the availability of backup origins.

**Figure 7.**
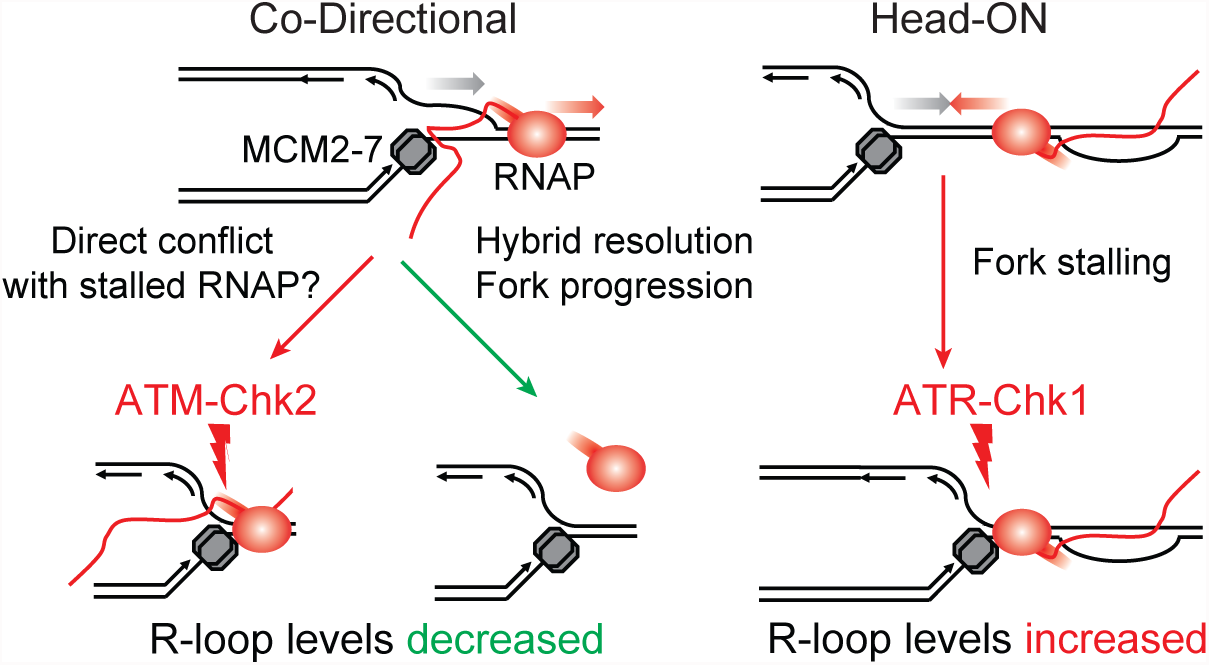
Model for HO and CD collisions in human cells. Head-on and co-directional transcription-replication conflicts regulate R-loop homeostasis and induce distinct DNA damage responses in human cells.

Our experimental approach provides intriguing insights into the impact of conflict orientation on TRCs. We show that HO collisions promote R-loop formation using our plasmid system and in the native genomic context (Figure 5D). We also find that HO collisions promote plasmid loss and ATR activation. Moreover, the instability of the R-loop forming mAIRN HO plasmid is partially reversed by RNaseH overexpression (Figure 3B), suggesting that the R-loop itself strongly contributes to genome instability in the HO orientation. We propose that R-loop forming transcription complexes are particularly strong blocks to replication fork progression in this orientation, consistent with robust phosphorylation of ATR and several of its downstream targets (Figure S6A). The stalled replication forks may lead to plasmid loss through a failure to complete replication and ultimately replication fork collapse. In bacteria, it has been debated whether fork stalling following HO collisions is a result of a direct clash between transcription and replication machineries or the accumulation of torsional stress that impairs progression even without physical contact (Hamperl and Cimprich, 2016; Mirkin et al., 2006). Although we cannot distinguish between these models and both factors may contribute in our plasmid system, the positive PLA signal between RNAP II and PCNA in this orientation (Figure 4B) supports the idea that the transcription and replication machineries are in close proximity on at least a fraction of the HO plasmids.

The precise mechanism for ATR activation following a HO collision is unclear. Replication-dependent ATR activation is known to occur as a result of ssDNA formed upon polymerase stalling and uncoupling of helicase and polymerase activities (Byun et al., 2005). However, in the context of a TRC, continued helicase unwinding may be blocked by the opposing RNAP complex, preventing ssDNA formation on the leading strand. We speculate that ssDNA is found on the lagging strand template, as part of the stalled replication fork, or is found within the R-loop, as part of the transcriptional barrier. The question also arises as to whether the increased formation of hybrids in the HO orientation is a cause or consequence of the collision event. The R-loop/transcription complex itself may cause replication fork stalling. Alternatively, the block to RNAP progression may keep the nascent RNA strand in proximity to the complementary DNA duplex for longer and thereby promote the formation of RNA-DNA hybrids as a consequence of the TRC. In fact, we observed a significant decrease of RNA-DNA hybrid levels in G1-arrested mAIRN HO cells compared to asynchronously growing or S-phase cells (Figure 2B). This finding suggests that R-loops are stabilized by the HO encounter with replication forks. Importantly, however, few hybrids were observed in ECFP HO cells despite the fact that HO collisions occur on this construct. Thus, we suggest that robust formation of R-loops may require both a HO collision and an R-loop prone sequence.

Conversely, we show that replication in the CD orientation decreases hybrid levels using our plasmid system (Figure 1D), consistent with reduced hybrid formation in genomic regions predicted to have a CD bias (Figure 5D). This finding suggests that replication may be a mechanism for R-loop resolution. An important question is whether there is a replisome-associated factor that executes this biochemical activity. Indeed, several candidate proteins with hybrid resolving activity have been found at stalled replication forks and could contribute to this process (Alzu et al., 2012; García-Rubio et al., 2015; Schwab et al., 2015; Yüce and West, 2013). However, another intriguing possibility is that the MCM2-7 helicase itself may directly resolve RNA-DNA hybrids in the CD orientation. This is consistent with the biochemical activity of related replicative helicases from all domains of life, which unwind RNA-DNA hybrids in 3’ to 5’ orientation (Shin and Kelman, 2006). It is also supported by the fact that the eukaryotic MCM helicase moves along the leading strand (Fu et al., 2011), which forms the hybrid in the CD orientation (Figure 7).

Despite the fact that hybrid levels decline in the CD orientation, we still observe plasmid instability and robust autophosphorylation of ATM when an R-loop is present, particularly at high levels of expression. This suggests that ATM is activated, consistent with the preferential phosphorylation of other ATM targets including Chk2 and KAP1 S824. Given that H2AX is a substrate of both ATM and ATR kinases (Podhorecka et al., 2010), the poor γ-H2AX signal in the CD orientation is surprising. We speculate that CD collisions may form a unique structure that does not promote efficient ATM-dependent H2AX phosphorylation or that there is a lack of H2AX molecules in the highly transcribed mAIRN CD sequence. In fact, γ-H2AX modification is strongly diminished over highly transcribed genes (Lee et al., 2014), and we achieved much higher transcription levels with the mAIRN CD cells than with the mAIRN HO cells (Figure 1C).

More importantly, the activation of ATM suggests that DSBs may be formed in the CD orientation, and this damage could account for plasmid instability in this orientation. One mechanism by which DSBs form may involve collision of the replication fork with a more stable form of the transcription machinery induced as a result of hybrid formation. In fact, paused or arrested RNAP complexes can backtrack along the DNA template, resulting in a highly stable but transcriptionally inactive conformation. Furthermore, these backtracked complexes have been shown to induce DSBs on bacterial plasmid templates in the CD orientation (Dutta et al., 2011). Thus, the backtracked polymerase resulting from hybrid formation may be a barrier that can cause fork arrest, DSB formation and ATM activation, despite the fact that the RNA-DNA hybrid may be resolved behind RNAP as the fork progresses. It is also possible that the TC-NER endonucleases XPG and XPF play a role in DSB formation, as these nucleases were recently implicated in processing unscheduled RNA-DNA hybrids into DSBs (Sollier et al., 2014; Stork et al., 2016). Further studies will be necessary to determine whether these or related flap-endonucleases, RNAP backtracking or another unknown mechanism causes DSB formation and consequent ATM activation in the context of CD-TRCs.

Importantly, our finding that R-loops are resolved by DNA replication in the CD orientation not only uncovers a novel intrinsic function of the replisome, but also provides new insight into how R-loops can fulfill their physiological functions without impairing DNA replication. Clearance of transient, regulatory R-loops with passage of the replication fork in the CD orientation would allow the cell to tolerate these structures with minimal impact on genome stability. By extension, this mechanism for R-loop resolution could also be important for the clearance of persistent R-loops that evade other R-loop resolution pathways and that threaten genome stability. Because replication forks must access the entire genome, this could be a safeguard that allows resolution of R-loops in genomic regions inaccessible to other R-loop resolution factors. Replisome-mediated hybrid clearance could also be important at replication forks stalled in the HO orientation, where the approach of another replication fork from the opposite orientation could allow resolution from the CD orientation.

Finally, our data show that replication slow-down or deregulation of origin firing induces hybrid accumulation by decreasing CD collisions and/or increasing HO collisions. Thus, proper execution of the replication program can suppress the accumulation of R-loops. We propose that this replication-dependent control of hybrid levels has important implications for our understanding of replication-stressed induced genome instability. Upon slowing or stalling of DNA replication forks, inactive or ‘dormant’ replication origins in the vicinity of the stalled fork are activated to complete DNA synthesis (Yekezare et al., 2013). Under conditions of replication stress, a hallmark of cancer cells induced by oncogene activation or nucleotide depletion (Bartkova et al., 2006; Bester et al., 2011), stalled replication forks accumulate inducing the activation of dormant origins. We propose that this elevation of dormant origin firing in a genome biased toward CD collisions may come at the cost of HO TRCs and increased occurrence of potentially genome-destabilizing R-loops, possibly saturating other R-loop resolution pathways. The co-orientation bias of the human genome may therefore help to coordinate replication with transcription, minimize deleterious R-loops, and maintain genomic stability.

Here, we focus on the effects of R-loops on TRCs and show that R-loops pose a greater threat to genome stability than transcription itself. Whether or not R-loops are unique in their ability to augment the deleterious impact of a TRC is an intriguing question in the field. Other events that impact transcription, such as transcription pause sites, may have similar effect. The episomal system we developed can be modified to address this and other questions about the impact and outcome of TRCs in mammalian cells.

## Author Contributions

S.H., J.C.S. and K.A.C. designed experiments. S.H. and J.C.S. performed experiments. M.B. performed the bioinformatical and statistical analyses. S.H., J.C.S. and M.B. analyzed the data. S.H. and K.A.C. wrote the manuscript.

## Acknowledgements

We thank Fréderic Chédin for reagents and protocols for *in vitro* transcription assays, Tomek Swigut, Fréderic Chédin and members of the Cimprich laboratory for helpful discussions, and Joanna Wysocka and Dan Jarosz for helpful feedback on the manuscript. This work was supported by grants from the NIH (currently, GM119334 and previously, GM100489) to K.A.C. J.C.S was supported by a Postdoctoral Fellowship, PF-15-165-01 - DMC from the American Cancer Society and holds a Postdoctoral Enrichment Program Award from the Burroughs Wellcome Fund. S.H. was supported by a fellowship from the German Research Foundation DFG (HA 6996/1-1).

## STAR Methods

### Cell Culture

Human embryonic kidney (HEK) cells were used as previous studies showed efficient replication of oriP/EBNA1 plasmids in this cell line (Leight and Sugden, 2001). HEK293 Tet-ON (Clontech) and HeLa cells (ATCC) were cultured in DMEM (GIBCO) supplemented with 10% FBS, 2 mM L-glutamine and penicillin/streptomycin in 5% CO2 at 37°C. To generate monoclonal HEK293-TetON cell lines, cells were transfected with vectors pSH26 (ECFP-HO), pSH27 (ECFP-CD), pSH36 (mAIRN-HO) or pSH37 (mAIRN-CD) and selected with 200 μg/ml hygromycin for 2-3 weeks. Surviving single cell colonies were further expanded and screened for stable maintenance and replication of the episomal DNA by quantitative PCR and Southern blotting. Cells were maintained under selection in 200 μg/ml hygromycin. For cell cycle synchronization experiments, cells were cultured in DMEM supplemented with 2mM thymidine for 19h, washed twice with PBS and released into S-phase for 6h with regular DMEM.

### Antibodies, RNA interference, and Reagents

Antibodies to γ-H2AX (Cell Signaling, 9718S), P-ATR S428 (Cell Signaling, 2853S), P-ATM S1981 (Cell Signaling, 4526S), P-CHK1 S345 (Cell Signaling, 2348S), CHK1 (Santa Cruz, clone G-4, sc-8408), P-CHK2 T68 (Cell Signaling, 2661S), CHK2 (Santa Cruz, sc-56297), P-KAP1 (Bethyl, A300-767A), KAP1 (Transduction Lab, K57620), PRPA32 S33 (Bethyl Laboratories, A300-246A), RPA32 (EMD-Millipore, clone Ab-3, NA19L-100UG), RNAP II (EMD Millipore, clone 8WG16, 05-952), pSer2 RNAP II (Abcam, ab24758), PCNA (Santa Cruz, sc-7907), MCM2 (Cell Signaling, clone 1E7, 12079S), MCM3 (Abcam, ab4460), BrdU (BD, 347580), ALPHA-TUB (Sigma, T9026), GAPDH (Abcam, ab8245) and FLAG (Sigma, F1804) are commercially available. The S9.6 antibody was purified from the S9.6 hybridoma cell line from ATCC. Hybridoma supernatant was applied to a 1 mL HiTrap Protein G HP column (GE Healthcare). The antibody was eluted with 100 mM glycine pH 2.5, in 0.5 mL fractions. Fractions were screened for antibody by SDS-PAGE and antibody-containing fractions were pooled and dialyzed in PBS overnight followed by dialysis in 50% glycerol for 6h. The antibody concentration was measured against a BSA standard and aliquots were made at 1 mg/ml. siRNAs, purchased from ThermoFisher, were: siGL3 (D-001400-01-20), siMCM2_1 (custom siRNA with the sequence 5’ GGA GCU CAU UGG AGA UGG CAU GGA A), siMCM2_2 (J-003273-09-0002), siMCM3_1 (custom siRNA with the sequence 5’ GCA UUG UCA CUA AAU GUU CUC UAG U), siMCM3_2 (J-003274-10-0002), siSRSF1 (D-018672-02-0002), siBRCA2 (L-003462-00-0005). All siRNA transfections were performed at 20nM in antibiotic free DMEM with 10% FBS using Dharmafect1 (ThermoFisher) according to the manufacturer’s protocol. Plasmid transfections were performed using Fugene 6 transfection reagent (Promega) following the manufacturer’s protocol. Doxycycline (DOX) (Sigma), hydroxyurea (Sigma), aphidicolin (Sigma) and Cdc7 inhibitor (PHA-767491) were added at the indicated concentrations and times.

### Oligonucleotides and plasmids

Unless noted otherwise, standard techniques were used for cloning of plasmids. Complete lists of oligonucleotides and plasmids used in this study can be found in Tables 1 and 2.

**Table 1.**
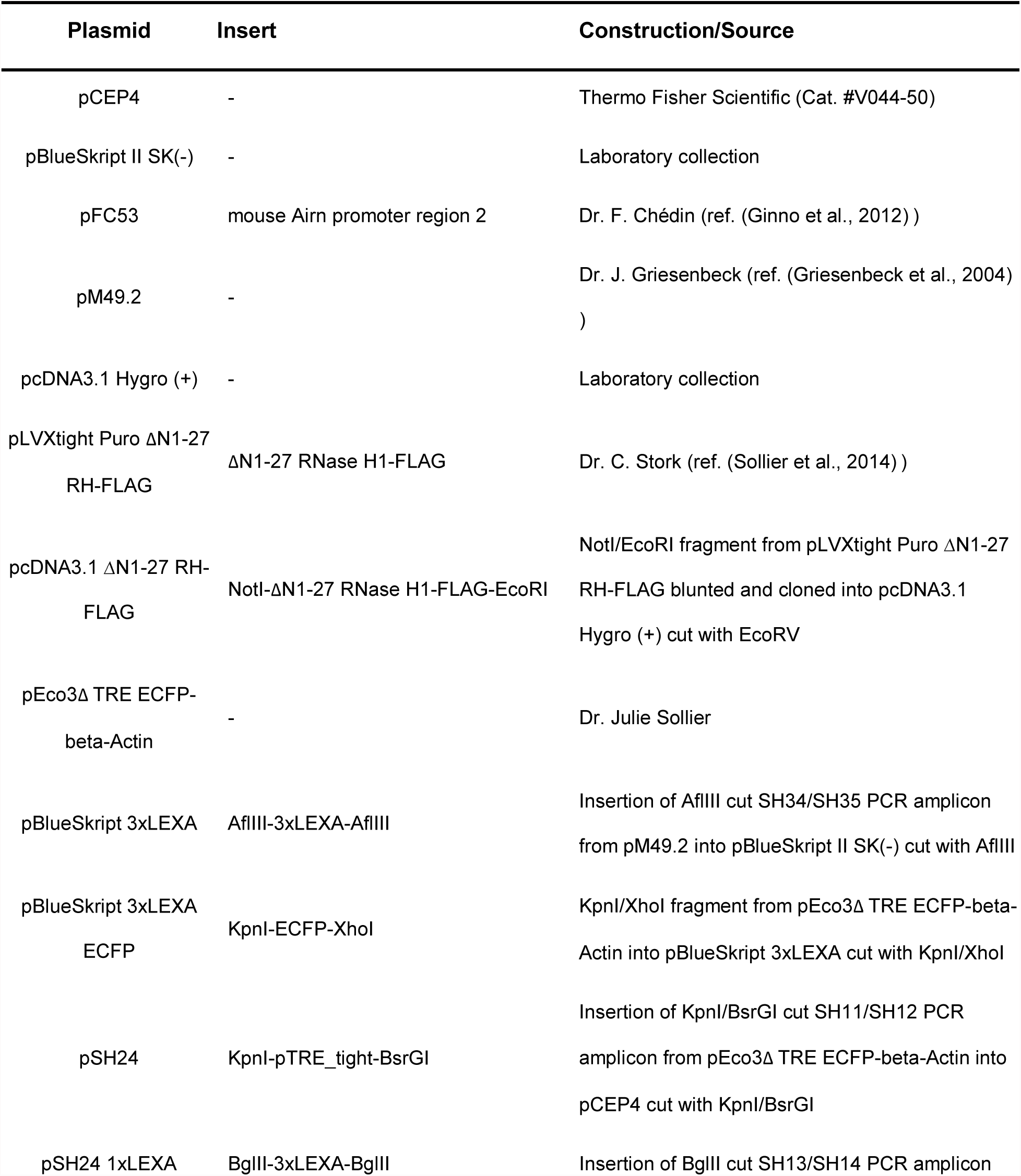
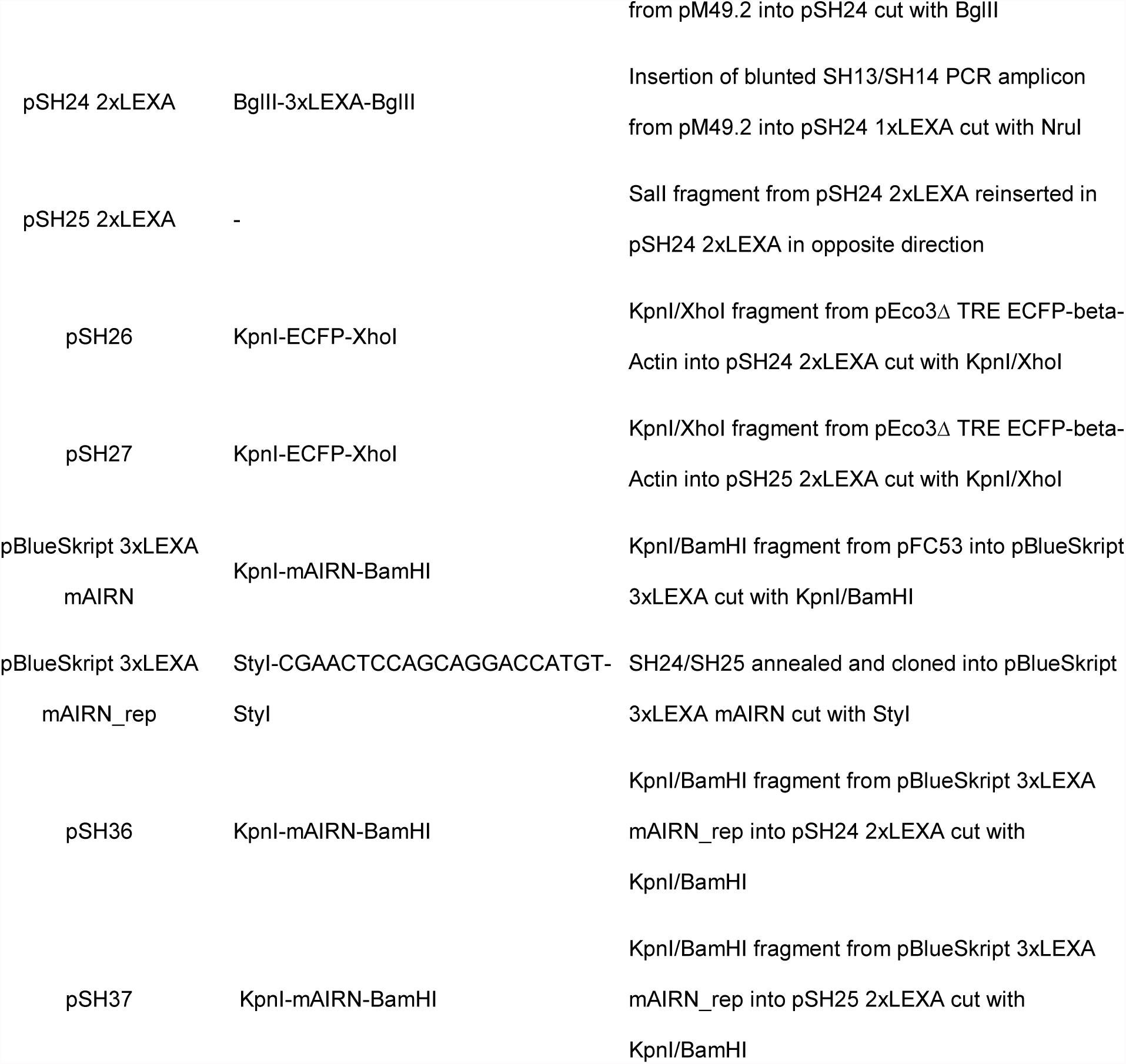
Table of plasmids used.

**Table 2.**
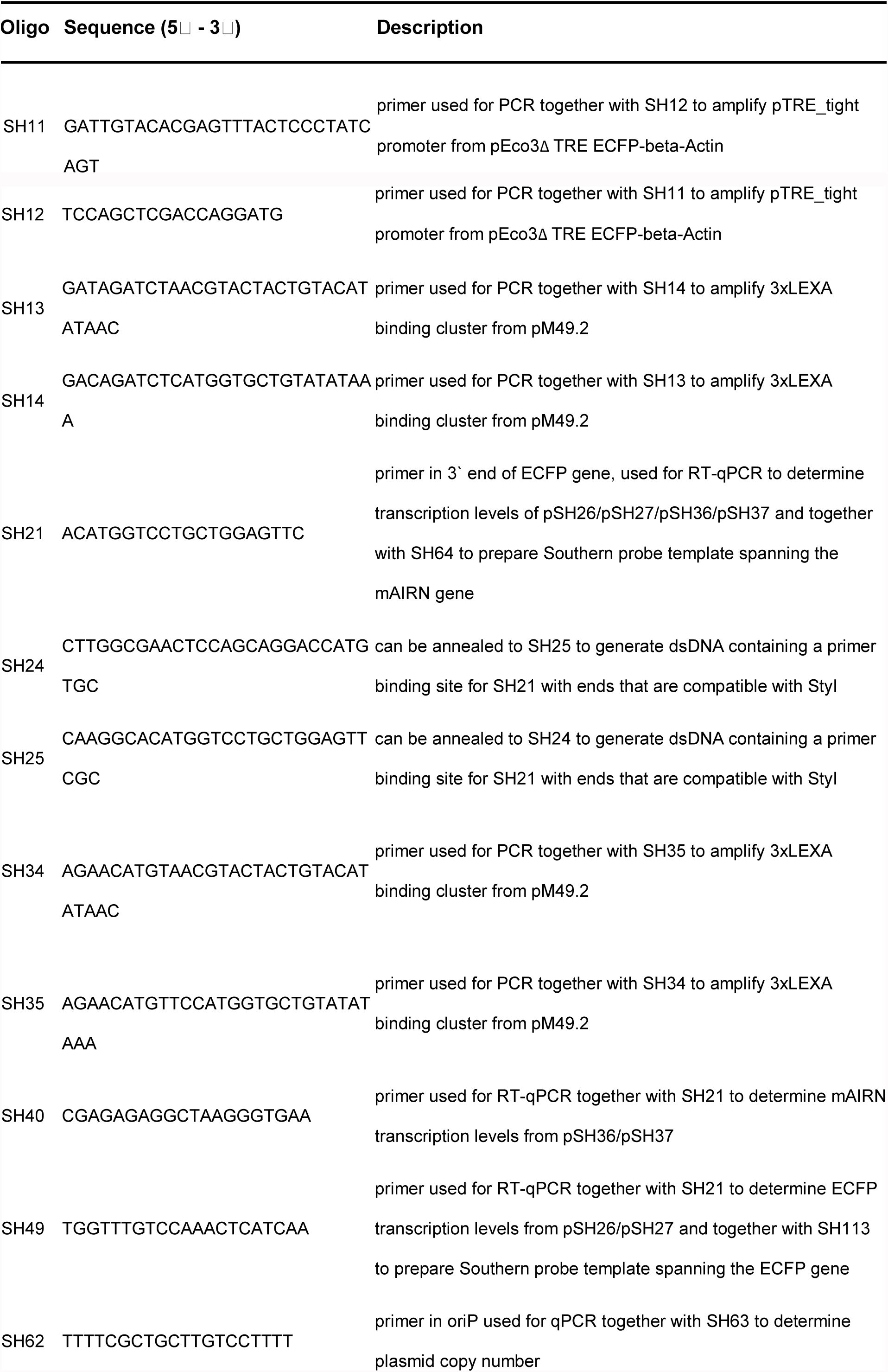
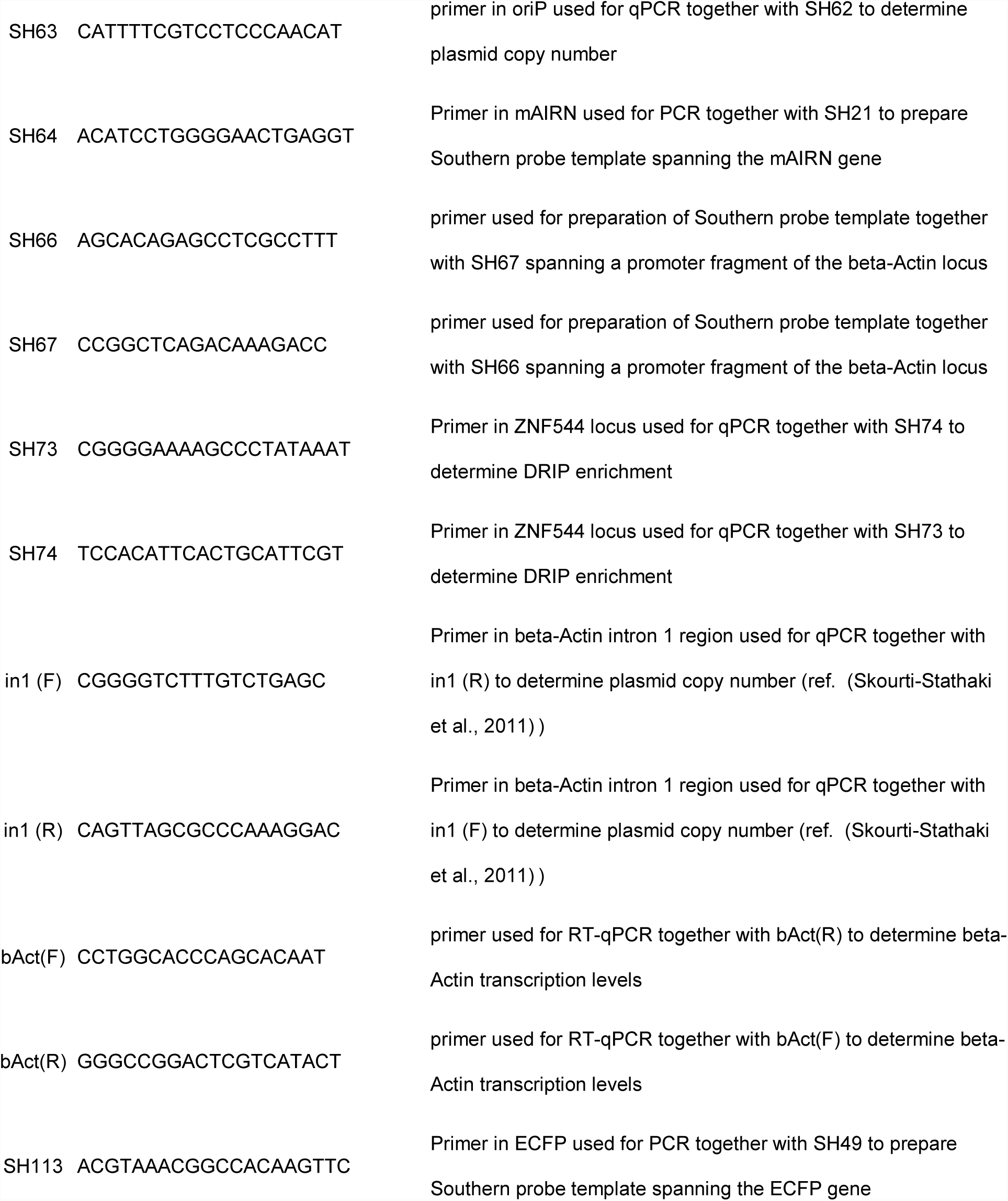
Table of oligonucleotides used.

### In vitro transcription for R-loop formation

Plasmid substrates (3 μg) containing the mAIRN or ECFP sequences flanked by oppositely oriented T3 and T7 promoters were *in vitro* transcribed using T3 or T7 RNA polymerase (Promega) according to manufacturer’s protocol at 37 °C for 30 min. After heat inactivation at 65°C for 10 min, the sample was split in half and incubated with 0.5 μg RNase A alone or 0.5 μg RNase A and 10U RNase H (NEB, M0297) as a negative control for 30 min at 37°C. After digestion with 20 μg Proteinase K for 30 min at 37°C, R-loop formation causes a characteristic shift in mobility of the plasmid on a 0.9% 1x TBE agarose gel run at 90V for 60 min. Gel was post-stained with ethidium bromide.

### Reverse Transcription-qPCR

Cells were harvested and total RNA was isolated using TRIzol reagent (Invitrogen) following the manufacturer’s protocol. After digestion with RNAse-free DNAse I (NEB, M0303) at 37°C for 30min, reverse transcription was carried out with 1.5 μg total RNA with random hexamer primers and SuperScript III Reverse Transcriptase Kit (Invitrogen). Equal amounts of cDNA were mixed with iTaq SYBR Green Supermix (Bio-Rad) and run on a Roche LightCycler 480 Instrument II. mRNA expression levels were measured by the change in comparative threshold cycles with primers in the beta-actin as a control.

### DNA:RNA immunoprecipitation

DRIP was performed as described in Ginno et al., 2012. Briefly, DNA was extracted with phenol/chloroform in phase lock tubes (5Prime), precipitated with EtOH/sodium acetate, washed with 70% EtOH, and resuspended in TE. DNA was digested with EcoRI and XcmI (NEB) restriction enzymes overnight at 37 °C. For RNase H-treated samples, 4 ug of DNA was treated with RNase H (NEB, M0297S) overnight at 37 °C. DNA was purified by phenol/chloroform, EtOH/sodium acetate precipitation as described above. 4μg of DNA was bound with 10 μg of S9.6 antibody in 1 X binding buffer (10 mM NaPO4 pH 7, 140 mM NaCl, 0.05% Triton X-100) overnight at 4°C. Protein A/G sepharose beads (Pierce) were added for 2 h. Bound beads were washed 3 times in binding buffer and elution was performed in elution buffer (50 mM Tris pH 8, 10 mM EDTA, 0.5% SDS, Proteinase K) for 45 min at 55°C. DNA was purified as described. Quantitative PCR of immunoprecipitated DNA fragments was performed on a Roche LightCycler 480 Instrument II using SYBR-Green master mix (Biorad).

### Plasmid copy number

Genomic DNA was isolated from 2-4 x 10^5^ cells. After trypsinization, cells were washed in 1x PBS and resuspended in TE buffer followed by the addition of an equal volume of IRN buffer (50mM Tris-HCl at pH 8, 20 mM EDTA, 0.5 M NaCl), 0.5% SDS and 10μg Proteinase K. After digestion for 1h at 37°C, DNA was extracted with phenol/chloroform and digested with 20μg RNase A for 1h at 37°C. After chloroform extraction, DNA was precipitated with EtOH/sodium acetate, washed with 70% EtOH, and resuspended in TE. DNA was digested with EcoRI (NEB) restriction enzyme overnight at 37°C. The plasmid copy number was analyzed with primer pairs SH62/SH63 and in1(F)/in1(R) amplifying either the oriP region of the plasmid or a region of the genomic beta-actin gene. The relative plasmid copy number was determined by quantitative PCR on a Roche LightCycler 480 Instrument II using SYBR-Green master mix (Biorad) and defined as the ratio of the amount of oriP to the amount of beta-Actin.

### Southern blotting

Southern blotting of genomic DNA was performed as previously described (Merz et al., 2008). Images were acquired with the Typhoon 9410 imaging system.

### Immunostaining

Cells were fixed with 4% PFA/PBS (EMS) for 15 min, permeabilized with 0.25% Triton-X 100 for 5 min, washed 3 times in 1X PBS, and blocked in 2% BSA/PBS for 1 hr at RT. For S9.6 immunofluorescence, cells were fixed in 100% methanol for 10 min at -20°C, washed 3 times in 1X PBS, and blocked in 2% BSA/PBS for 1 hr at RT. Primary antibody was incubated overnight at 4°C. Antibodies: Rabbit P-H2AX antibody (1:500, Cell Signaling), P-ATR S428 (Cell Signaling, 1:100), P-ATM S1981 (Cell Signaling, 1:100), S9.6 (1:100), Nucleolin (Abcam, 1:500). Cells were then washed 3 times in 1X PBS and co-stained with Hoechst (1:1000) and anti-rabbit AlexaFluoro-488- and anti-mouse AlexaFluoro-594-conjugated secondary antibodies (1:1000). Cells were imaged at 63x on a Zeiss AxioObserver Z.1 Fluoresence Light Microscope or Zeiss LSM 500 Confocal Microscope with ZEN 2009 Software. Analysis of P-H2AX, P-ATR or P-ATM intensity per nucleus was calculated using Image J (v 1.50b), where Hoechst is used as a mask for the nucleus. In box and whisker plots, box and whiskers indicate 25-75 and 10-90 percentiles, respectively, with lines representing median values.

### Cell Cycle Analysis

To monitor S-phase progression, cells were pulse-labeled with 25 μM 5-Bromo-2′- deoxyuridine (BrdU) for 30 min, and washed three times with PBS. After fixing samples with ice-cold 70% ethanol, cells were permeabilized with 0.25% Triton X-100/PBS for 15 min on ice, blocked in 2% BSA/PBS for 15 min, and incubated in primary BrdU antibody (BD Bioscience) for 2 h. Cells were then washed three times in PBS, incubated in AlexaFluoro-488 secondary antibody for 1 h, and washed three times with PBS. Propidium iodide (PI; 0.1 mg/mL; Sigma) and RNase A (10 mg/mL) was added to determine DNA content and cells were analyzed on a FACSCalibur device (BD Bioscience). Cell cycle profiles were determined using FlowJo™ software.

### Proximity Ligation Assay

For the proximity ligation assay (PLA), cells were pre-extracted with cold 0.5% NP-40 for 4 min on ice. Cells were then fixed with 4% PFA/PBS for 15 min, washed 3 times with 1X PBS and blocked for 1 h at RT with 2% BSA/PBS. Cells were then incubated in primary antibody overnight at 4°C (1:500 mouse RNAP II 8WG16Pol; 1:500 rabbit PCNA alone; or 1:500 mouse RNAP II 8WG16 with 1:500 rabbit PCNA). Cells were then washed 3 times in 1X PBS and incubated in a pre-mixed solution of PLA probe anti-mouse minus and PLA probe anti-rabbit plus (Sigma) for 1 h at 37°C. The Duolink In Situ Detection Reagents (Green) were then used to perform the PLA reaction according to the manufacturer’s instructions. Slides were mounted in Duolink In Situ Mounting Medium with DAPI and imaged on a Zeiss Axioscope at 40X or on a AxioObserver Z.1 at 63x. The number of PLA foci was quantified using Image J.

### DRIP-Seq

HeLa cells were cultured in DMEM (GIBCO) supplemented with 10% FBS, 2 mM Lglutamine and penicillin/streptomycin in 5% CO2 at 37°C and transfected with siGL3 for 72h using Dharmafect1 (ThermoFisher) according to the manufacturer’s protocol. DRIP followed by library preparation, next generation sequencing, and peak calling were performed as described in Stork et al., 2016 with minor modifications. RNase A pretreatment before DRIP was conducted with 6μg/ml of enzyme for 45 min at 37°C in 10 mM Tris-HCl pH 7.5 supplemented with 0.5M NaCl as described in Sanz et al., 2016. In addition, sheared Drosophila melanogaster chromatin (Active Motif, 53083) and a Drosophila-specific H2Av antibody (Active Motif, 61686) was spiked-in as a minor fraction of the DRIP DNA to allow normalization of read counts according to the manufacturer’s protocol. Libraries were pooled and sequenced on an Illumina HiSeq 4000 machine with paired-end 75bp reads at the Stanford Functional Genomics Facility (NIH grant S10OD018220). The raw sequencing data were uploaded to the GEO database (access number pending).

### Bioinformatic and statistical analyses

Replication fork directionality (RFD) in HeLa cells was obtained from the OK-Seq data in the supplementary information of (Petryk et al., 2016). Using the mean RFD between the two replicates, the data was smoothed using a cubic spline fit. Origin-proximal areas were defined as all areas of the spline with positive slope between a local minimum and a local maximum. These areas were split into a Crick region (stretching from the local minimum to the discrete zero) and a Watson region (stretching from the discrete zero to the local maximum). In order to classify the local transcriptional profile around these origins, we used GRO-seq data in HeLa cells (GEO dataset GSM1518913). Using bedtools (Quinlan and Hall, 2010), we calculated the total GRO-seq read density over the Watson and Crick portions of each origin-proximal area, for both positive and negative stranded transcription. Using the relative direction of transcription in each area, we classified each origin as either HO-HO, CD-CD, HO-CD, or CD-HO. The last two categories were condensed into a single class of origins, by inverting the genome coordinates for any CD-HO origins. With these classified origins, we calculated profiles of DRIP-seq read density in a 30kb window around each type of origin using the HTseq python package and matrix manipulations in numpy (Anders et al., 2015). In order to account for different levels of confounding variables known to correlate with DRIP signal, we calculated the mean replication timing (using HeLa RepliSeq profiles from ENCODE), transcription levels (using the previously mentioned GRO-seq data) and GC-content (using bedtools) across each 30kb region. We matched the categories of origins to each other by standardizing these variables, then finding the closest (not necessarily unique) element in the match set to each element in the test set, using the L2 distance on the three match variables. Using these matched pairs between the samples, we then bootstrapped the test sample 50,000 times. For each round of bootstrapping, we used the matching algorithm to find a comparable sample in the matched sample. Using these bootstrapped and matched samples, we plotted the mean and 95% confidence interval of the mean for each sample.

---

**Figure S1.**
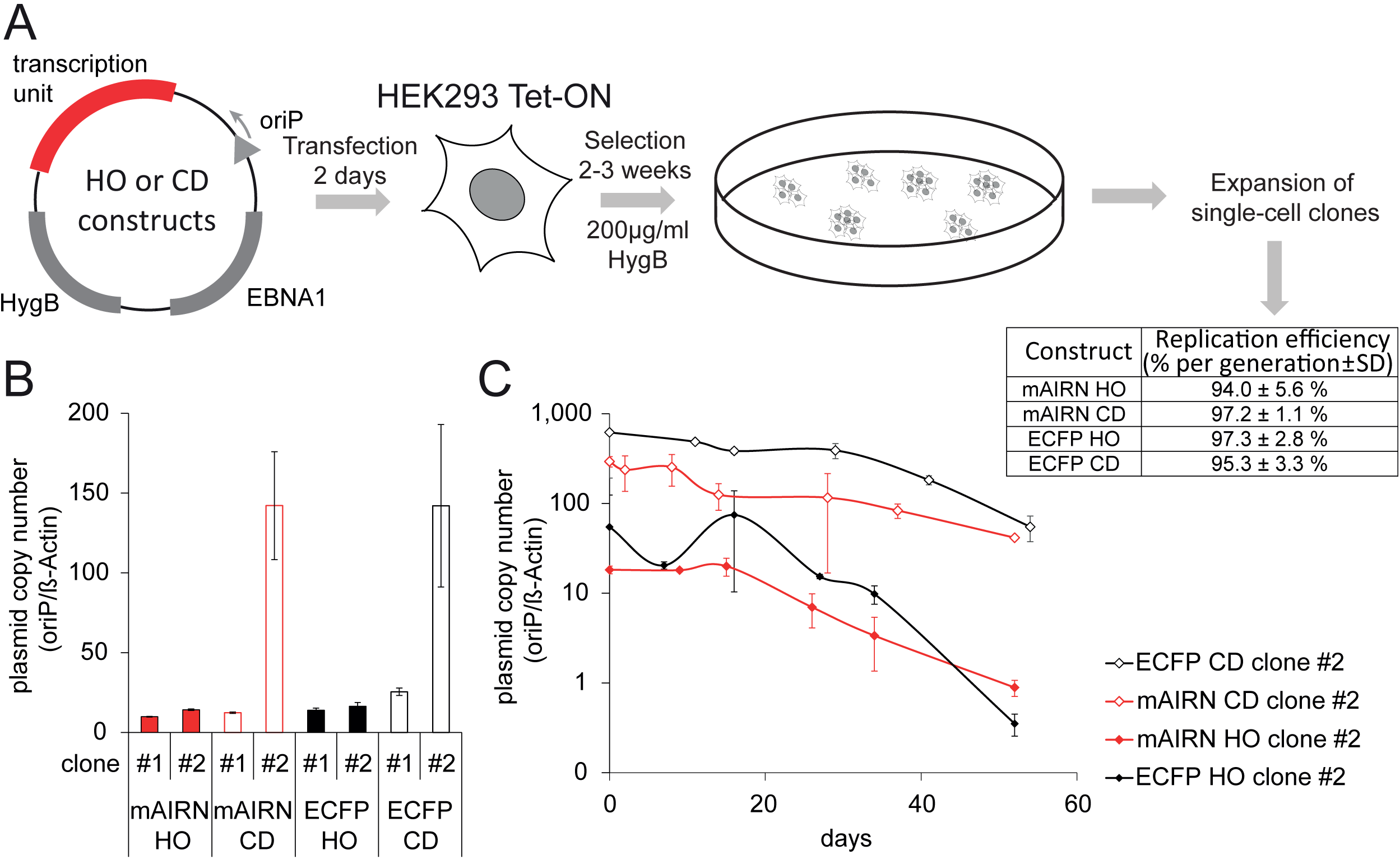
Isolation and characterization of single cell clones that efficiently replicate the mAIRN and ECFP constructs. A) The oriP plasmid constructs expressing EBNA1 and hygromycin phosphotransferase (HygB) were transfected into HEK293 Tet-ON cells. Two days after transfection, 10^2^ to 10^3^ cells per 10-cm dish were plated in media containing 200 μg/ml hygromycin B. After selection for 2 to 3 weeks, ~20-40 drug-resistant clones for each plasmid derivative were isolated and expanded, giving rise to the mAIRN and ECFP HO and CD cell clones used in this study. B) The plasmid copy number of the individual cell clones was determined by qPCR analysis after the initial expansion of the clones. If not otherwise stated, clone #2 cells for each construct were used for analyses throughout this study. C) Replication efficiency of each construct was monitored over the course of ~8 weeks by qPCR analysis of the plasmid copy number remaining in the cell population. The bars indicate mean and standard deviations between biological replicates (n=3).

**Figure S2.**
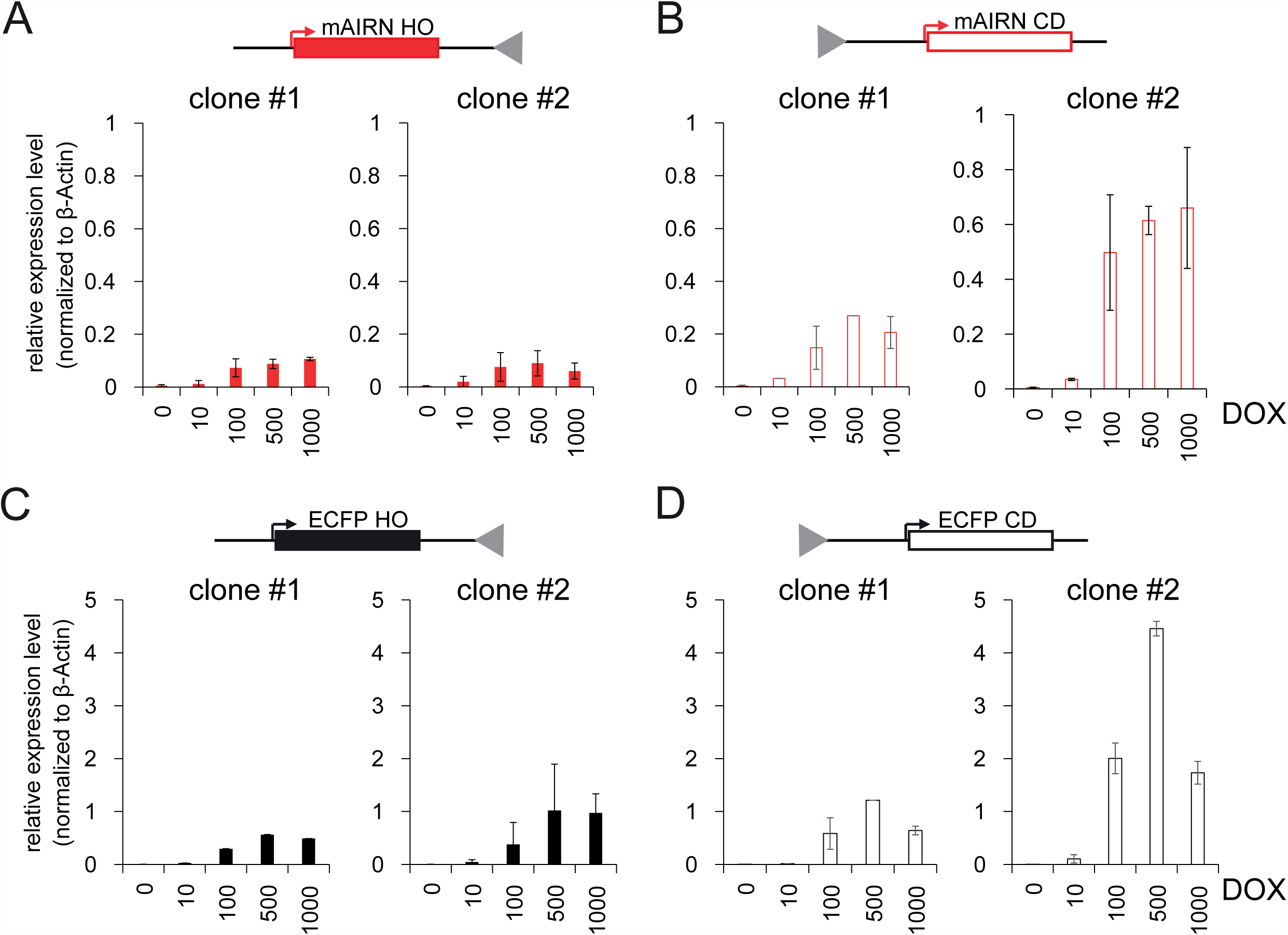
mAIRN and ECFP expression in co-directional and head-on orientations. A-D) RT-qPCR analyses of RNA samples extracted from cells induced with 0, 10, 100, 500 or 1000 ng/mL DOX for 72h. Gene expression was measured and normalized relative to β-Actin as a reference gene. The bars indicate mean and standard deviations between biological replicates (n=3, except for 10 and 500 ng/mL DOX for mAIRN CD clone #1 and ECFP CD clone #1 where n=1).

**Figure S3.**
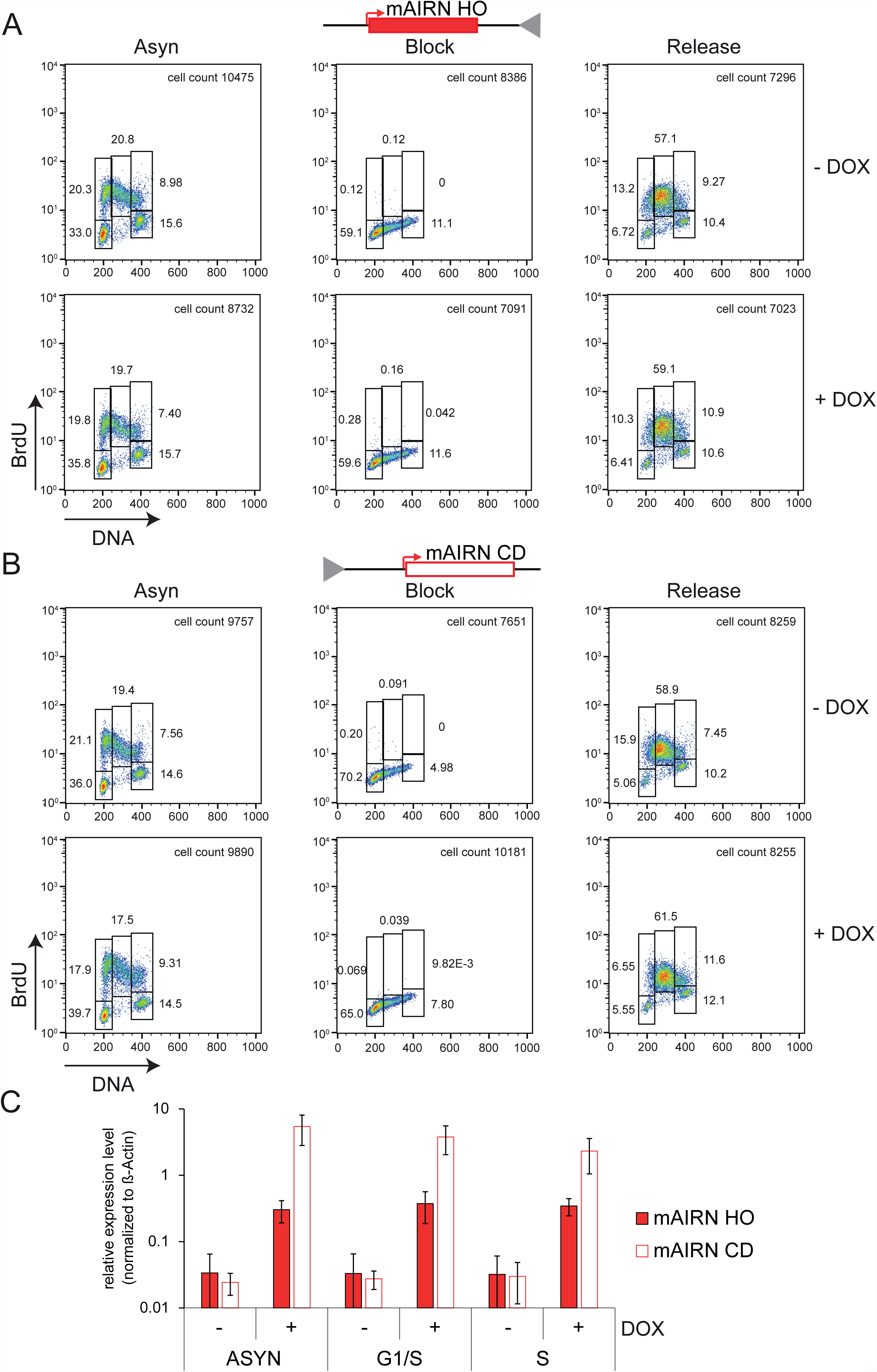
Flow cytometry and RT-qPCR analyses after synchronization of mAIRN HO and CD cells with thymidine. A-B) Representative fluorescence-activated cell sorting (FACS) profiles of mAIRN HO (clone #2) and CD (clone #2) cells treated with 0 (-) or 1000 ng/mL (+) DOX under asynchronous conditions (ASYN), after treatment with 2mM thymidine for 19h (Block) or subsequent wash-out with fresh medium for 6h (Release). Cells were pulsed with 25 μM BrdU for 30 min prior to fixation. DNA content is marked by propidium iodide as shown on the x-axis and BrdU incorporation is shown on the y-axis. The percentage of cells in G1, early, mid and late S and G2/M-phase is shown. C) RT-qPCR analysis of mAIRN HO and CD cells under the conditions described in A) and B). RNA samples were extracted and gene expression was normalized relative to the expression of the β-actin gene. The bars indicate mean and standard deviations between biological replicates (n=3).

**Figure S4.**
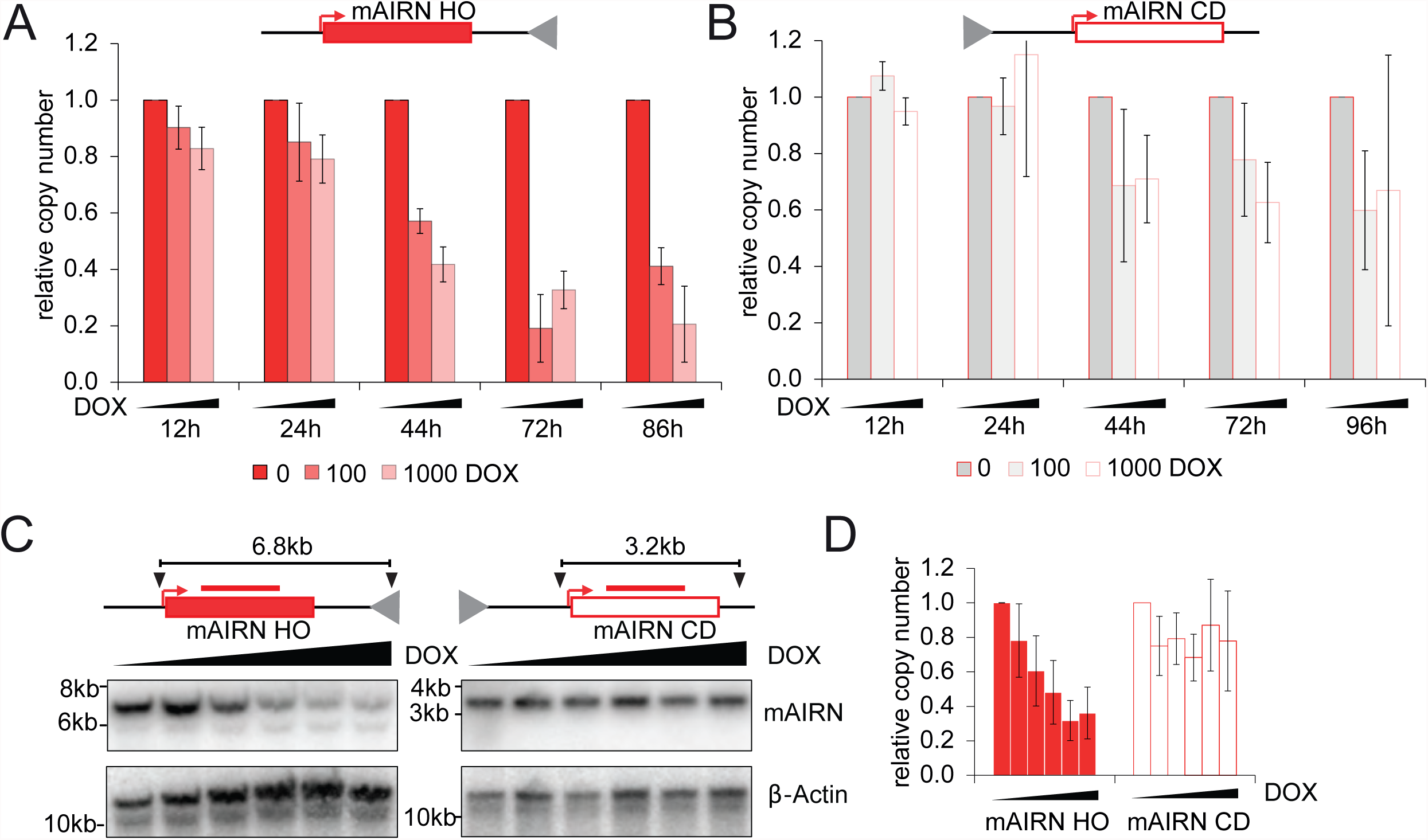
Transcription-induced plasmid instability is orientation and R-loop dependent. A-B) mAIRN HO cells (clone #2) or mAIRN CD cells (clone #2) were treated with 0, 100 or 1000 ng/mL DOX for the indicated timepoints. After extraction of genomic DNA, the relative plasmid copy number (normalized to 0ng/ml DOX) was determined by quantitative PCR. The bars indicate mean and standard deviations between biological replicates (n=3). C) Representative Southern blot of EcoRI digested DNA samples from mAIRN HO or mAIRN CD cells after induction of transcription with 0, 10, 50, 100, 500 or 1000 ng/mL DOX for 72h. Black triangles and red bars indicate positions of EcoRI restriction sites and the mAIRN probe used to visualize a 6.8kb or 3.2kb fragment of the mAIRN HO or CD construct, respectively. D) Quantification of Southern blot experiments as shown in C). The relative plasmid copy number was determined by the ratio change of the mAIRN fragment and an EcoRI fragment derived from the genomic β-actin locus (normalized to 0ng/ml DOX). The bars indicate mean and standard deviations between biological replicates (n=2).

**Figure S5.**
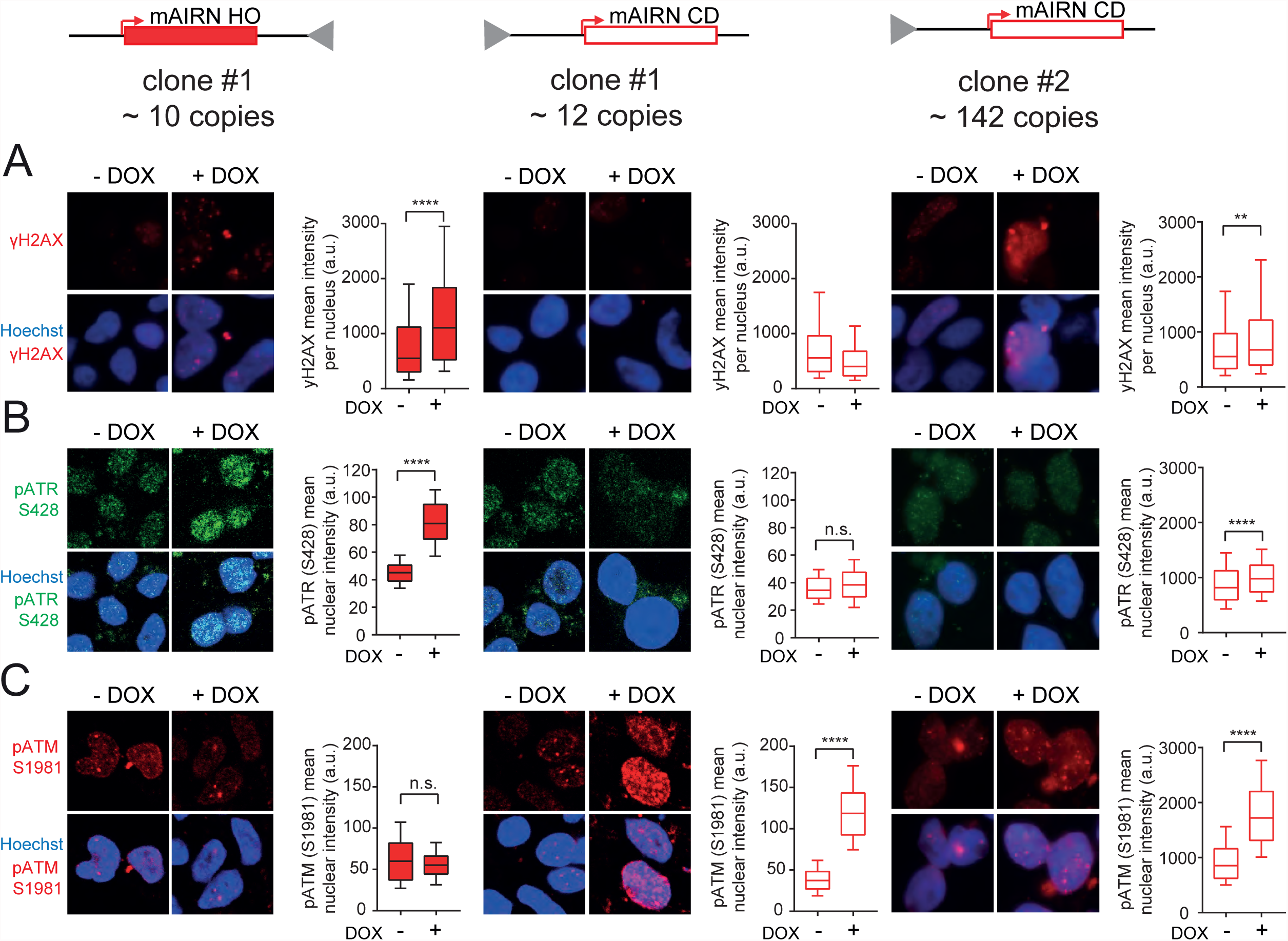
Preferential activation of ATR or ATM checkpoint kinases in mAIRN HO and CD cells. A-C) Representative images (left) and quantification (right) after immunostaining for A) γH2AX, B) ATR pS428 or C) ATM pS1981 in mAIRN HO (clone #1) and mAIRN CD (clone #1 or clone #2) cells either treated with 0 (-) or 1000 ng/mL (+) DOX for 48h. Hoechst is used to stain the nucleus. Box and whisker plots show the 10-90 percentile. a.u. = arbitrary units. n.s. not significant. **p<0.01. ****p<0.0001. One-way ANOVA test (n ≥ 120).

**Figure S6.**
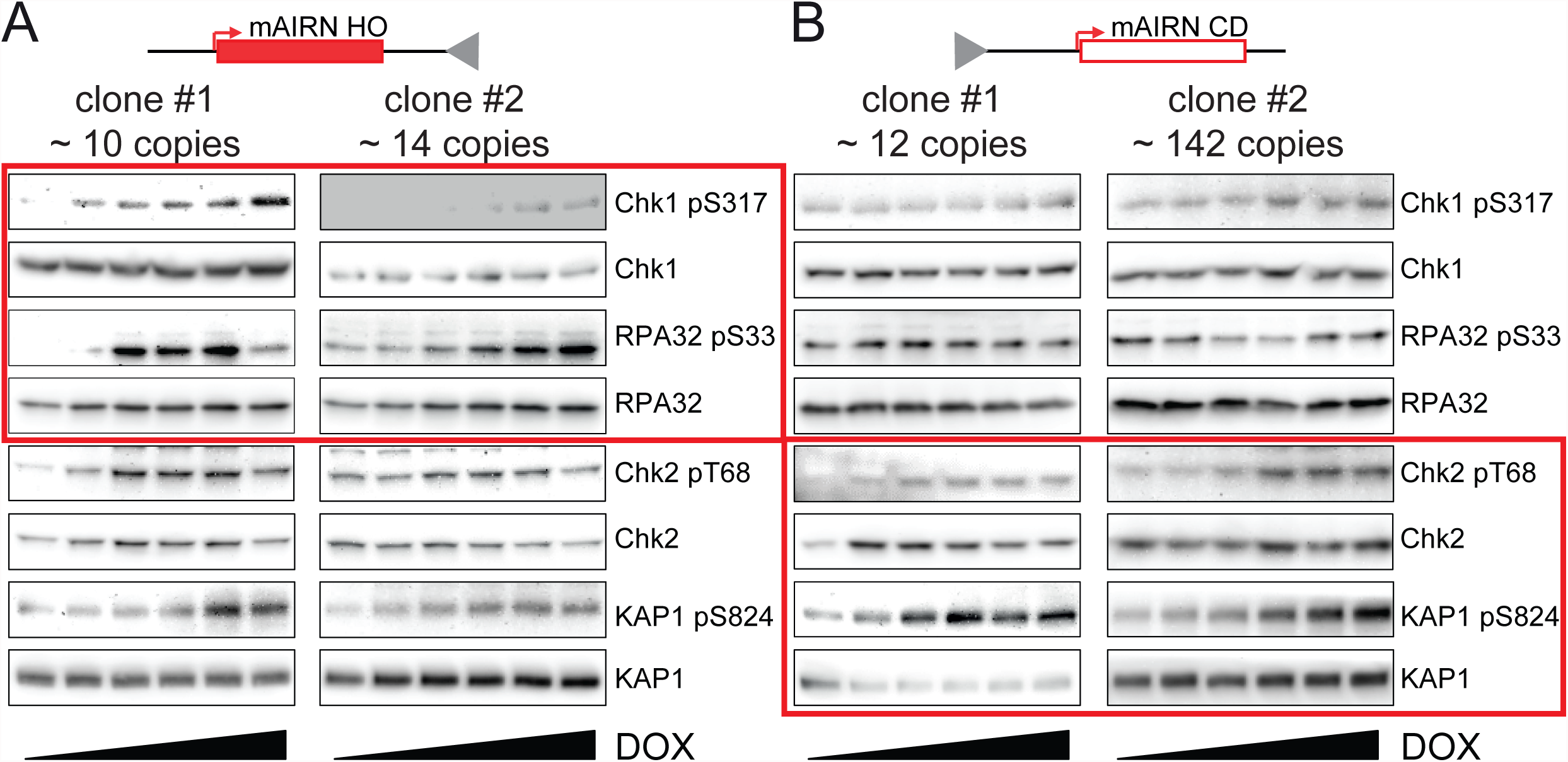
Western blot validation of the specific DNA damage checkpoint responses in mAIRN HO and CD cells. A-B) mAIRN HO or mAIRN CD cells were treated with 0, 10, 50, 100, 500 or 1000 ng/mL DOX for 48h. After preparation of whole cell lysates, equal protein amounts were analyzed by immunoblotting with the indicated antibodies.

**Figure S7.**
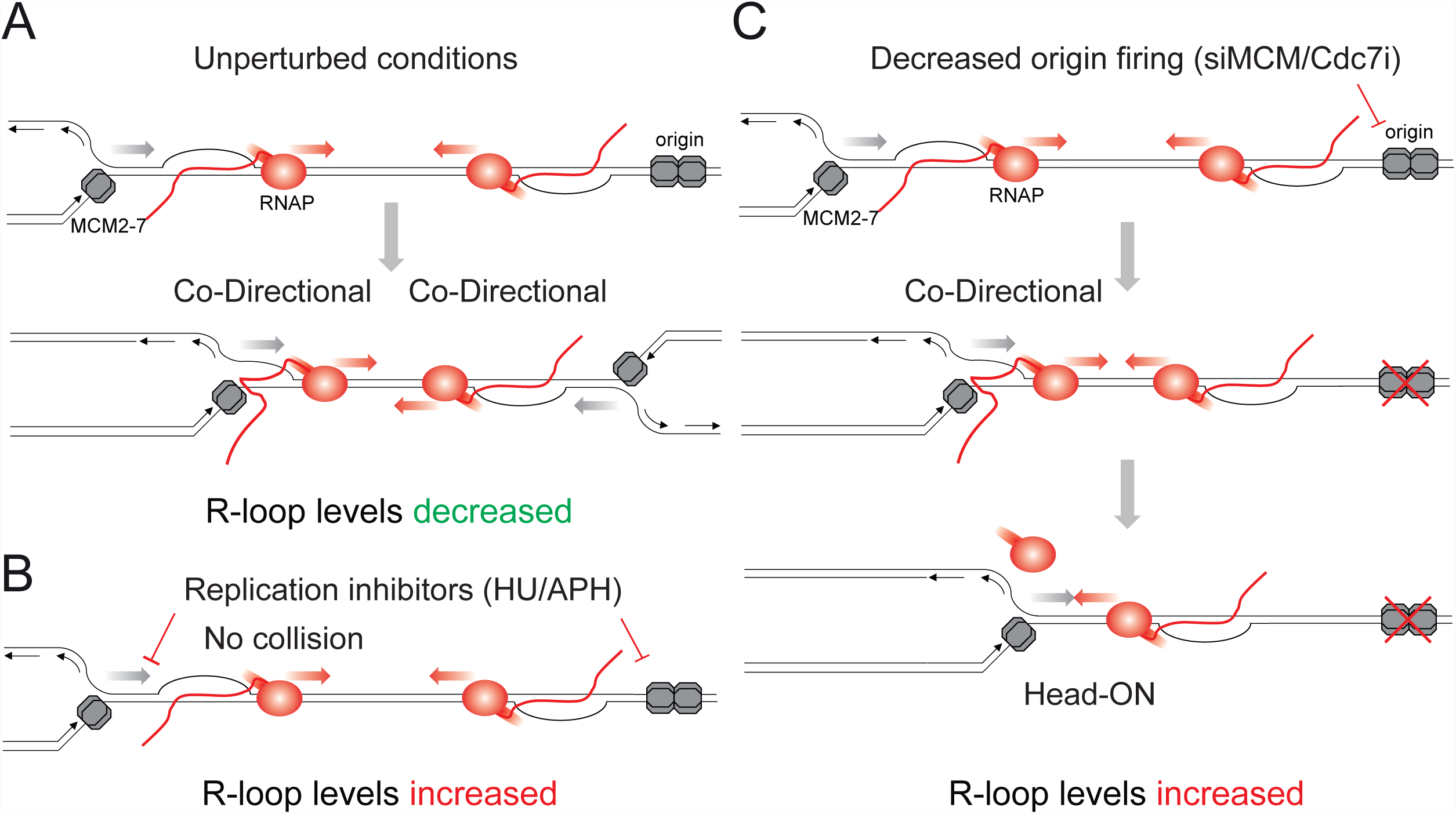
Model illustrating how different perturbations of the replication program regulate R-loop homeostasis in human cells. A) Under unperturbed conditions, R-loop levels are decreased by the preferential co-directional movement of replication forks and transcription complexes. This bias towards CD collisions allows the replisome to resolve RNA-DNA hybrids, most likely by processive CMG (Cdc45-Mcm2-7-GINS) helicases that encircle the RNA-DNA hybrid containing leading strand in the CD orientation to drive replication fork progression (Fu et al., 2011). B) Replication inhibitors (HU/APH) induce R-loops by blocking replication fork progression, thereby inhibiting the resolution of RNA-DNA hybrids during CD collisions. C) Decreasing the number of active origins by depletion of Mcm proteins or inhibition of Cdc7 kinase reduces the number of active replication forks and/or changes the pattern of origin usage, therefore increasing the frequency of HO collisions with transcription complexes and increased R-loop levels.

**Figure S8.**
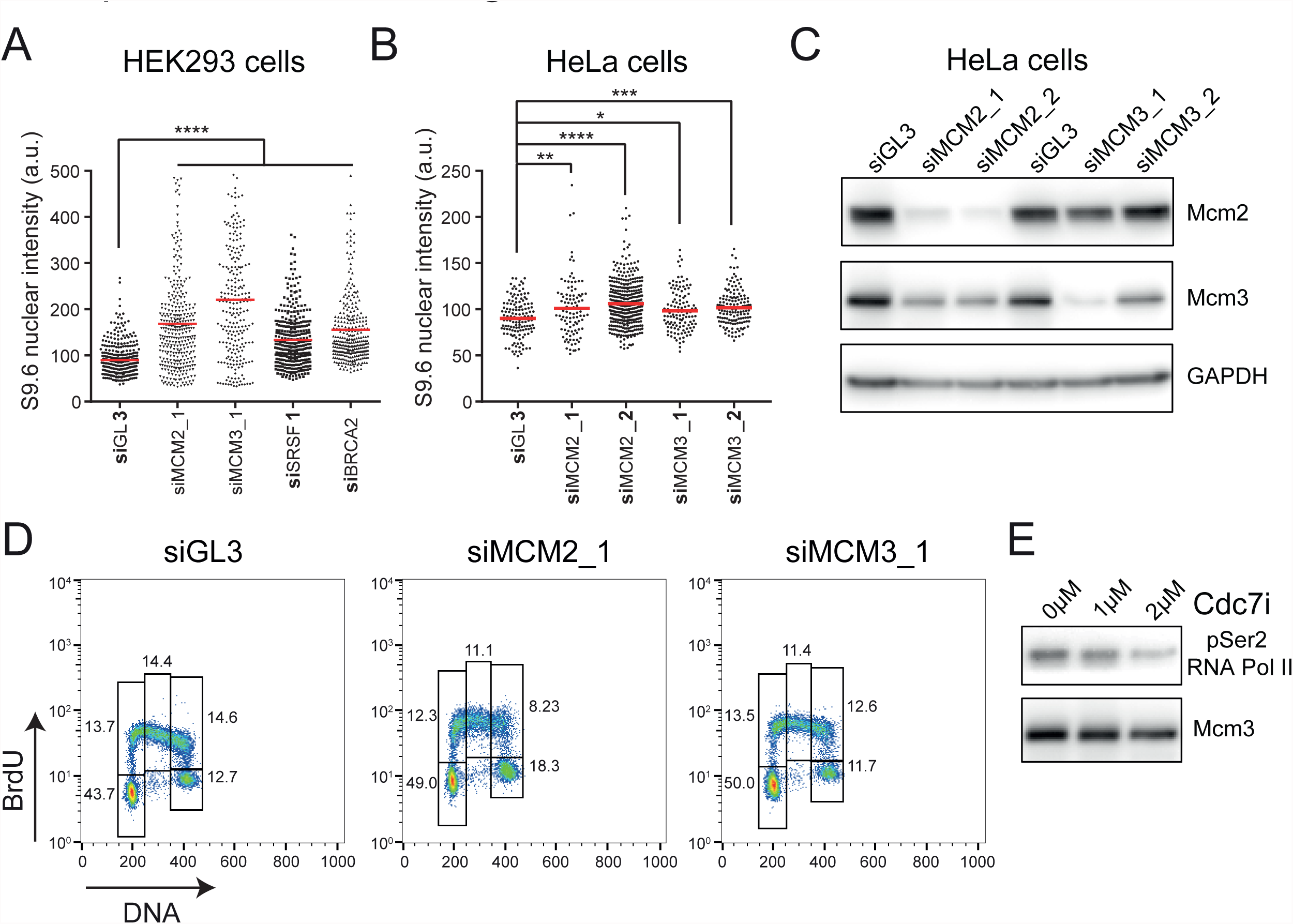
Specificity of siRNAs and Cdc7 inhibitor and cell cycle distribution after MCM depletion. A) Quantification of S9.6 nuclear signal in HEK293 cells transfected with indicated siRNAs and fixed after 72h. The nucleus was co-stained with Hoechst. The mean value is shown as a red line. a.u. = arbitrary units. ****p<0.0001. One-way ANOVA test (n≥100). B) Quantification of S9.6 nuclear signal in HeLa cells transfected with indicated siRNAs and fixed after 72h. The nucleus was co-stained with Hoechst. The mean value is shown as a red line. a.u. = arbitrary units. *p<0.05. **p<0.01. ***p<0.001. ****p<0.0001. One-way ANOVA test (n≥100). C) Western blot analysis of HeLa cells transfected with indicated siRNAs. After 72h, whole cell lysates were analyzed by immunoblotting with the indicated antibodies. D) Representative FACS profiles of HeLa cells transfected with indicated siRNAs as in B). Cell cycle profiles were acquired as described in the legend to Figure S3A-B. E) Western blot analysis of HeLa cells treated with Cdc7 inhibitor (PHA-767491) at the indicated concentrations. After 4h, whole cell lysates were analyzed by immunoblotting with the indicated antibodies.

